# A Standardized Method for Insect Color Analyses using Open Source Software: AInsectID Version 1.1 Color Merge

**DOI:** 10.1101/2025.05.28.656714

**Authors:** Haleema Sadia, Sahal Sabilil Muttaqin, Parvez Alam

## Abstract

The accurate representation of color is important in applications involving species identification. Environmental variations introduce inconsistencies in color perception, affecting the reliability of automated image processing algorithms. In previous work, we developed a hybrid algorithm, AInsectID Version 1.1 Color Merge, to overcome challenges posed by over-segmentation and under-segmentation in insect wing color clustering. We achieved this by using color differences between superpixels to measure homogeneity during the superpixels segmentation process. Nevertheless, our algorithm remains sensitive to environmental effects, affecting its performance and accuracy in color analyses. Here, we introduce a standard imaging method for insect species, as a pre-requisite to analysis in AInsectID Version 1.1 Color Merge. We systematically examine the effects of varying lighting conditions, angle of observation, and working distance in a controlled environment to assess their impact on the performance of the algorithm. We find that by meticulously controlling lighting, working distance, and lighting angle, we develop an evidence-based standard approach to imaging colors that is robust and repeatable. By following our standardized procedure, consistent color analyses are possible under varying environmental conditions. The method was tested using the Delta E2000 (Δ*E*) color difference metric with a threshold of 1, demonstrating that our standard approach maintains perceptual accuracy within the Just Noticeable Difference (JND) range, while improving the reliability of color analyses of insect wings in diverse environments. Finally, to validate the robustness of our standardization method, we evaluated the certainty of our results at different levels of confidence.

## 1 Introduction

In nature, coloration arises from the intrinsic properties of materials, alongside physical phenomena such as light diffraction and interference. This phenomenon, known as structural coloration or iridescence, has long fascinated scientists [1–7]. Studies have shown that these colors are created by intricate micro- and nanostructures, fundamental optical mechanisms including thin-film and multilayer interference, diffraction gratings, and three-dimensional photonic crystals [2] [8]. The coloration of insect wing scales largely depends on the manipulation of light during the interaction [4], light scattering, and the angle of observation [1]. Despite advancements in modern nanotechnology, replicating the phenomenon of natural colorization remains a significant challenge [9, 10].

In natural structures like insect wings [11], bird feathers [12], and chameleon skin [7], structural coloration is produced by photonic band gaps resulting from the Bragg scattering of light through periodic nanostructures, rather than relying solely on the chemical properties of pigments [6]. Natural organisms have evolved complex structures that combine high reflectivity at specific wavelengths with diffuse light distribution across broad angles, making color analysis difficult because of the intricate and dynamic nature of their coloration [11, 13]. Studies on Lycaenid species reveal that multilayered scales and asynchronous bending contribute to discoloration effects, thereby enhancing recognition [5]. Studies on *Heliconius sara* reveal that actin plays a critical role in scale development, with densely packed actin bundles forming reflective ridges responsible for iridescence. Actin perturbation experiments confirm its vital role, as disruptions result in a loss of structural color, suggesting the existence of a universal patterning mechanism in Lepidoptera [14]. The structural coloration in *Sternotomis virescens* longhorn beetles arises from amorphous photonic networks within elytral scales, producing green-blue patterns on black elytra. However, accurately analyzing these complex color effects remains a difficult task, due to the intricate nature of the photonic structures involved [15].

Machine learning (ML) and computer vision applications have demonstrated exceptional performance in insect identification and image processing [16–19], offering valuable tools to analyze morphology and color patterns [20–22]. However, these still face certain levels of uncertainty and unreliability due to the unique and complex structural properties of biomaterials [2–6, 8]. Modern computer vision technologies operate on the principle that objects emit or reflect light, which is captured by a camera’s pixels. The measured light intensities at each pixel are referred to as pixel values. Variations in light intensity, direction, and quality can alter how a camera perceives and records color, leading to differences in pixel values that may affect overall image quality and color accuracy [23].

Computer vision algorithms for image segmentation analyze pixel values to group similar regions called superpixels. Superpixels group pixels in an image to reduce the complexity of the image while retaining important visual information. Unlike individual pixels, superpixels represent homogeneous regions that are more consistent in color and texture, making them an ideal unit for image analysis tasks such as segmentation, recognition, and tracking. These algorithms rely on inherent patterns in the image, such as color, texture, or intensity, to determine segmentation boundaries. The reliability of the output from the algorithm has a high dependence on image quality, since all processing is based on pixel-level data [24, 25]. Segmentation results can fluctuate substantially under varying lighting conditions and angles of observation, as changes in illumination affect pixel values and disrupt detected patterns [26]. For example, shadows or uneven lighting may cause a wall in a room to appear as distinct segments rather than a single cohesive structure. This sensitivity to variations in lighting highlights a key limitation in current computer vision segmentation methods, leading to inconsistent results in real-world applications [27].

Standard clustering techniques, which typically assume uniform and isotropic cluster distributions, are often inadequate for managing color variability. When lighting effects are not sufficiently accounted for, segmentation algorithms may result in over-segmentation, where homogeneous regions become unnecessarily divided, or under-segmentation occurs, where subtle color distinctions left unnoticed [28] [29] [30]. These problems can lead to incomplete or erroneous representations of insect wing morphology [31]. In conclusion, machine learning algorithms continue to encounter significant challenges due to the intricate structural properties of biological materials, characterized by high reflectivity at specific wavelengths and diffuse light distribution across broad angles. These factors introduce uncertainties that affect the accuracy and consistency of computer vision-based algorithms, ultimately compromising the precision of wing morphology analyses [32]. Despite the numerous computer vision applications for color clustering [33–35], there is currently no standardized algorithm that performs consistently well across varying lighting conditions [36]. This lack of standardization remains a major obstacle in ensuring reliable and reproducible segmentation results.

In our previous work [26], we addressed the challenges of color clustering in insect wings by introducing a hybrid approach combining superpixel-based segmentation and color clustering techniques [37] incorporated into the AInsectID Version 1.1 Color Merge software from the AInsectID Version 1.1 package available in [38]. We utilized the Simple Linear Iterative Clustering (SLIC) method to generate the initial superpixels and employed the DeltaE 2000 function (Δ*E*) to discriminate and merge the superpixels precisely. This approach was designed to overcome the difficulties posed by the intricate color patterns of insect wings, which can lead to over- or under-segmentation when using traditional clustering algorithms. By measuring color differences between superpixels, our method ensured better homogeneity and more accurate segmentation, thereby improving the representation of the complex color diversity inherent in insect wing structures. It achieves better segmentation accuracy, as validated by metrics such as Boundary Recall, Rand Index, Under-segmentation Error, and Bhattacharyya distance. This paper extends its application by developing a formal standard for its use. This work aims to establish a set of guidelines that can be universally applied to ensure the algorithm’s reproducibility and adaptability under varying environmental and imaging conditions. By defining a standard for preprocessing steps, parameter selection, and output validation, we aim to provide a framework that guarantees consistent performance across different datasets, and imaging conditions.

## 2 Why is Standardization Needed?

The AInsectID Version 1.1 Color Merge algorithm is designed to perform precise segmentation of butterfly wing images by ensuring accurate color discrimination. It utilizes a superpixels colors merging threshold of 0.5, which adheres to the Just Noticeable Difference (JND) standard for color perception, quantifying the smallest detectable difference between two colors that a normal human eye can perceive. In image processing and display technologies, perceptual accuracy and color consistency are important [39]. If the calculated difference between two superpixel colors is less than 1 (in Δ*E* units), it is deemed indistinguishable from normal human vision [26]. The JND in color is commonly measured using the Δ*E* (Delta E) metric, which quantifies color differences, as established by the International Commission on Illumination (CIE). Generally, a Δ*E* value of approximately 1 represents the JND for most people, which means that differences below this value are almost imperceptible. A Δ*E* value between 1 and 2 may be noticeable under close observation, while values above 2 or 3 indicate a clearly perceptible difference. If Δ*E* exceeds 5, the color difference is considered obvious [40, 41]. While our algorithm merges superpixels precisely, its reliability is affected by variations within the input images taken under different lighting conditions and angles. Variations in lighting often result in inconsistent color merging, as the RGB values of merged colors frequently exceed the JND threshold (Δ*E*=1). This inconsistency is further conflated by the intricate structural coloration of insect wings, which reflect and refract light differently depending on the environment and observation angle. These factors lead to challenges in maintaining consistent performance under diverse environments, as illustrated in Figure 1. Varying lux (lumens per square meter) values affect the perception of color, causing noticeable shifts in the RGB values of butterfly wing coloration. These changes highlight the sensitivity of the AInsectID Version 1.1 Color Merge algorithm to lighting conditions, surpassing the JND threshold. As such, we consider standardization as being critical in the context of color merging and clustering, to ensure both reliability and consistency when using the AInsectID Version

**Figure 1.**
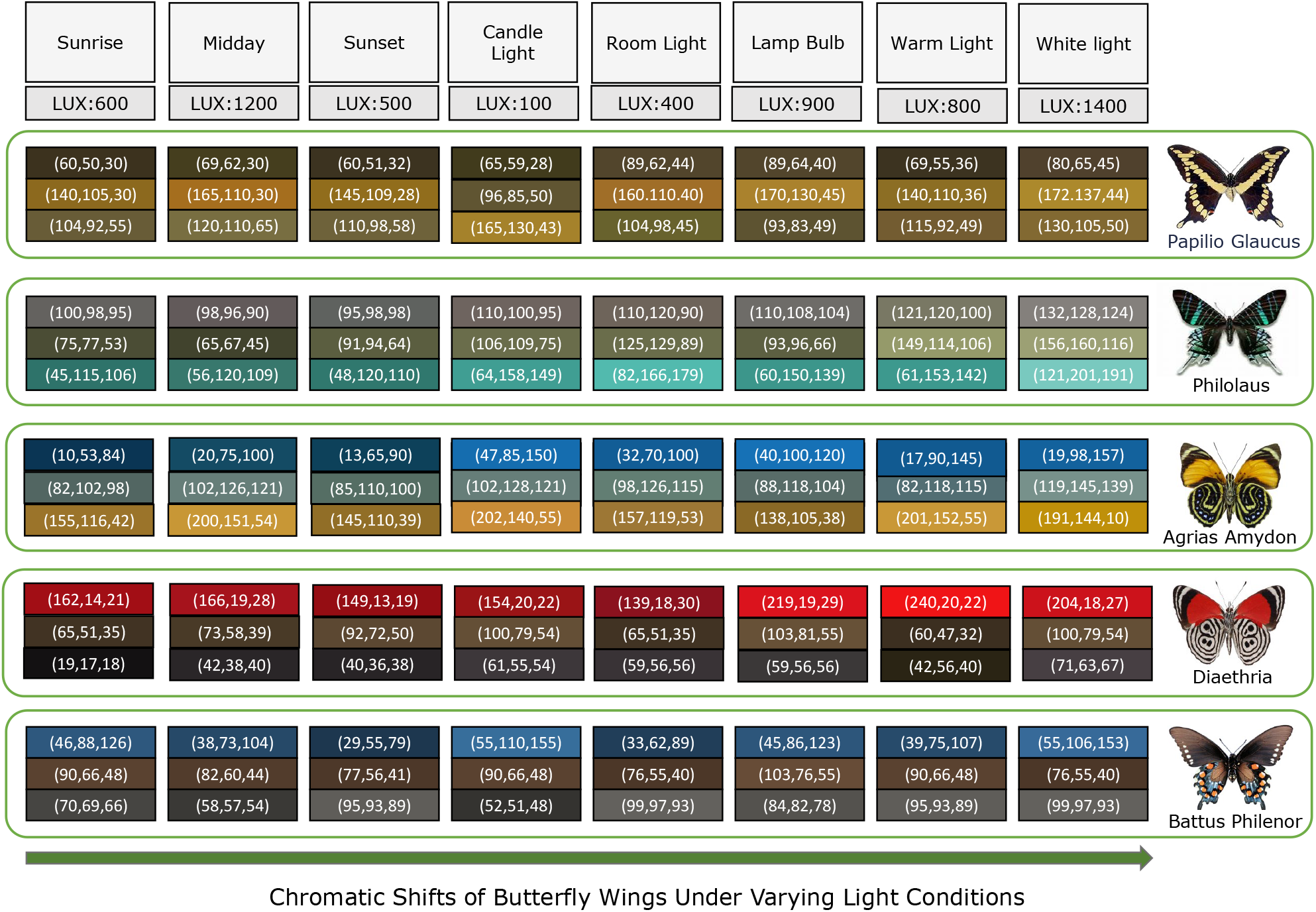
Comparison of RGB values of butterfly wing colors under varying lighting conditions, illustrating how the AInsectID Version 1.1 Color Merge algorithm processes and merges identical colors differently. These lighting variations lead to color shifts exceeding the Just Noticeable Difference (JND) threshold (Δ*E*=1), emphasizing the influence of lighting conditions on AInsectID Version 1.1 Color Merge algorithm consistency and reliability

1.1 Color Merge algorithm under varying lighting and angles of observation. By standardizing input data and controlling environmental variables, we can ensure consistent color rendering while maintaining perceptual accuracy. This will make the algorithm a reliable tool for both research and practical applications.

## 3 Entomological Photography Guidelines

In entomological photography, variations in sunlight can pose significant challenges, as it may create shadows and highlights that distort the true colors and details of the insect. Light intensity can vary reflectivity and if intense, can result in overexposure, particularly on glossy or iridescent surfaces. These factors make it difficult to capture the true appearance of an insect [42]. To mitigate these issues, entomology photographers often diffuse the sunlight and use controlled techniques to soften the light. One common approach is to use a reflector to redirect sunlight onto the insect from a more favorable angle, reducing shadows and evenly illuminating the subject [13]. In digital microscopic photography, ‘diffusers’ such as translucent materials or fabrics, are placed between the insect and the sun to soften the light and reduce glare, spreading the lighting more evenly and making it appear natural. Additionally, shade is used to shield the insect from direct sunlight, again to create a uniform lighting environment. In some cases, ‘fill flash’ is employed to add additional light to the subject without the creation of high reflectivity, especially when the sunlight casts deep shadows. By managing sunlight in this way, photographers can reduce the negative effects of intense lighting, ensuring that the insect is subjected to soft and even illumination, thereby enabling improved color reproduction, accurate detail capture, and overall higher image quality [42–44].

Entomologists use ‘white LEDs’ in controlled light environments to provide steady, adjustable illumination for insect photography. These lamps offer consistent light without heat or flicker, enabling them to capture accurate detail. Adjustable light intensity helps avoid deep shadowing and high reflectivity, especially on glossy surfaces. Paired with a white background, the setup enhances color accuracy and clarity, making it easier to discern insect morphology and achieve high-quality, reproducible images. Additionally, white LEDs cancel out the effect of external light, ensuring a consistent and controlled lighting environment. This controlled lighting setup is also ideal when capturing museum specimens, as it provides a consistent and precise illumination needed for detailed examination and accurate documentation in a variety of collections [45–49].

Insect photography commonly uses lenses with focal lengths from 50mm to 200mm. 50mm to 100mm lenses are affordable but require close proximity to the subject, which may disturb the insect. 100mm to 150mm lenses strike a balance, allowing detailed shots from a moderate distance with good background blur. 150mm to 200mm lenses offer greater distance, ideal for photographing skittish insects without disturbing them. Overall, 100mm to 150mm lenses are considered the best for most insect photography, offering a mix of sharpness, working distance, and versatility. Moreover, these lenses typically use 1:1 a magnification, or life-size magnification, where the subject appears at its actual size on the image sensor. This is ideal for the capture of finer details on insects in their natural state. For closer imaging, magnifications higher than 1:1, such as 2:1 or 5:1, can be used to show intricate details like wing texture or body hair. Nevertheless, a 1:1 magnification is usually sufficient for most insect photography, offering a good balance between detail and an ability to photograph the sebject without disturbance [48, 50, 51].

DSLR and mirrorless cameras are commonly used for photgraphing insects. This is due to their high resolution, versatility with interchangeable lenses, and ability to capture fine details using macro lenses. Popular models include the Canon EOS series, Nikon D series, and Sony Alpha series. These cameras provide excellent image quality and rapid autofocus, essential for capturing intricate features on insects bodies. Point-and-shoot cameras with macro capabilities, such as the Canon PowerShot and Nikon Coolpix series, are also used for their convenience and portability, especially in field settings. For highly detailed close-up shots, specialized cameras with high-resolution sensors, such as the Phase One XF IQ4 or Canon EOS 5DS R, are utilized and often paired with macro lenses to achieve 1:1 magnification. Macro lenses, extension tubes, and focus stacking techniques further enhance the ability to capture minute details, making them essential tools in entomological imaging [52]. Aperture selection is critical for depth of field and sharpness for macro photography. f/8 to f/11 is commonly used, balancing detail and focus for morphological analysis. For a greater depth of field, f/16 to f/22 can be employed, though diffraction can reduce sharpness, requiring controlled lighting such as diffused flashes. Wider apertures (f/5.6 to f/8) are used to enable selective focus, highlighting features like eyes or wings, while blurring the background. For extreme close-ups, focus stacking combines multiple images to achieve maximum detail. Aperture choice depends on the subject, lighting, and the desired scientific or artistic outcome [53, 54]. The 65mm f/2.8 1-5*×* macro lens is a specialized lens designed exclusively for macro photography of insects, offering a magnification of up to 5:1. Its compact and easy-to-handle design makes it perfect for capturing extreme macro shots, especially in situations where higher magnifications are required. To capture detailed images of insects, a shallow depth of field (DOF) is often recommended in order to isolate the subject from the background, while ensuring the insect remains the focal object of the photograph. The 100mm f/2.8 macro lens captures sharp, detailed images with effective background separation [55].

The 180mm f/2.8 macro lens is highly regarded for its exceptional image quality and optical stabilization, making it an excellent choice for macro photographers seeking precise focus and steady shots. In contrast, the 150mm f/2.8 EX DC OS HSM macro lens offers both optical stabilization and a hyper-sonic motor for quiet, fast, and accurate focusing, allowing photographers to capture insects and other delicate subjects without disturbing them. These lenses are all top contenders with an ability to produce outstanding insect macro photographs [56].

While considering the different entomological photography guidelines, we carefully designed our experimental setup to ensure the accurate capture of insect images under standardized conditions. This involved the implementation of a controlled lighting setup to minimize variabilities caused by external factors such as ambient light, shadows, and reflections. Once the images were captured under these standardized conditions, we tested the AInsectID Version 1.1 Color Merge algorithm to evaluate its performance. The aim was to ensure that the algorithm could consistently merge colors within the images without deviations caused by lighting inconsistencies. This approach allows us to standardize the input data for processing, ensuring consistency and reliability across different specimens and environmental conditions.

## 4 Research Methodology

To evaluate the algorithm’s robustness, we developed an experimental setup that systematically adjusted lighting conditions while capturing images of insect wings. The collected image data was processed using the AInsectID Version 1.1 Color Merge algorithm, allowing us to analyze how different lighting conditions, working distances, and angles of observation influence color consistency. By quantifying color differences across different setups, we established a standardized approach that ensures accurate and consistent color merging.

To objectively measure the differences in color, we employed the DeltaE 2000 (Δ*E*_00_) function with a threshold of 0.5, as shown in Equation (1). The Δ*E*_00_ formula is used to calculate color differences between two colors in the Commission on Illumination (CIE) L*a*b* color space, providing a more perceptually accurate measurement than previously reported [57]. This method accounts for sensitivities in human vision by incorporating factors such as lightness, chroma, and hue differences, along with perceptual non-uniformity corrections. To compute Δ*E*_00_, the colors are first converted into the *L*^*^*a*^*^*b*^*^ color space, where *L*^*^ represents lightness, *a*^*^ corresponds to the greenred axis, and *b*^*^ represents the blue-yellow axis. Several intermediate values are then computed, including weighted lightness difference (Δ*L*^*′*^), chroma difference (Δ*C*^*′*^), and hue difference (Δ*H*^*′*^), where *S*_*L*_, *S*_*C*_, *S*_*H*_ are weighting functions for lightness, chroma, and hue respectively, *k*_*L*_, *k*_*C*_, *k*_*H*_ are parametric weighting factors for lightness, chroma, and hue respectively, and *R*_*T*_ accounts for hue rotation effects that account for interactions between chroma and hue differences.

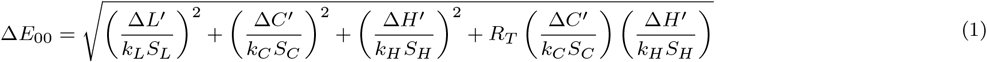

When Δ*E*_00_ ≤ 1, the colors are essentially indistinguishable to the human eye. This is often considered acceptable in industries such as textiles, paint, and digital imaging, where slight variations are permissible. However, when Δ*E*_00_ *>* 1, the colors are deemed perceptibly different, with larger values indicating a more noticeable difference. In quality control and color matching, differences greater than 1 may be undesirable, particularly in fields requiring precise color accuracy, such as paint manufacturing, graphic design, or fashion, where even minor deviations can impact consistency and visual appeal. In this study, we use the Δ*E*_00_ formula to calculate the color difference merged by the AInsectID Version 1.1 Color Merge algorithm to assess its reliability and consistency under varying environmental conditions. The threshold 0.5 ensures that the color differences remain virtually imperceptible to the human eye, which is essential for applications requiring high color accuracy [58]. This choice is particularly critical in contexts where even minimal color variations can impact visual consistency and product quality, ensuring that the color merge algorithm performs reliably and consistently under varying environmental conditions.

### 4.1 Experimental Setup

Following entomological imaging guidelines, we standardized the lighting conditions, angle of light relative to the insect wing, and of observation, working distance, and magnification for photography to ensure an accurate and consistent representation of color patterns. To ensure a controlled lighting environment, the experiment was conducted in a room with 0 ambient lux. As shown in Figure 2, a white light board was used as the background, four adjustable white light LED lamps were positioned to illuminate the specimen’s top surface to eliminate enhance color pattern visibility, a tripod stand used to ensure stable camera positioning, and a lux meter was used to measure and standardize the light intensity. Imaging was conducted using a DSLR camera Monitech4K UltraHD 48MP, featuring a macro lens with 16x digital optical zoom, 52mm, and an aperture of f/5.6, ensuring high-resolution capture of fine morphological details. The 52mm macro lens is particularly well-suited for close-up work with dead insects, as it allows us to focus at very short distances and achieve sharpness, and offers an optimal balance between 1:1 magnification and depth of field for detailed images of the specimens [59], making it an appropriate choice for the requirements of this study.

**Figure 2.**
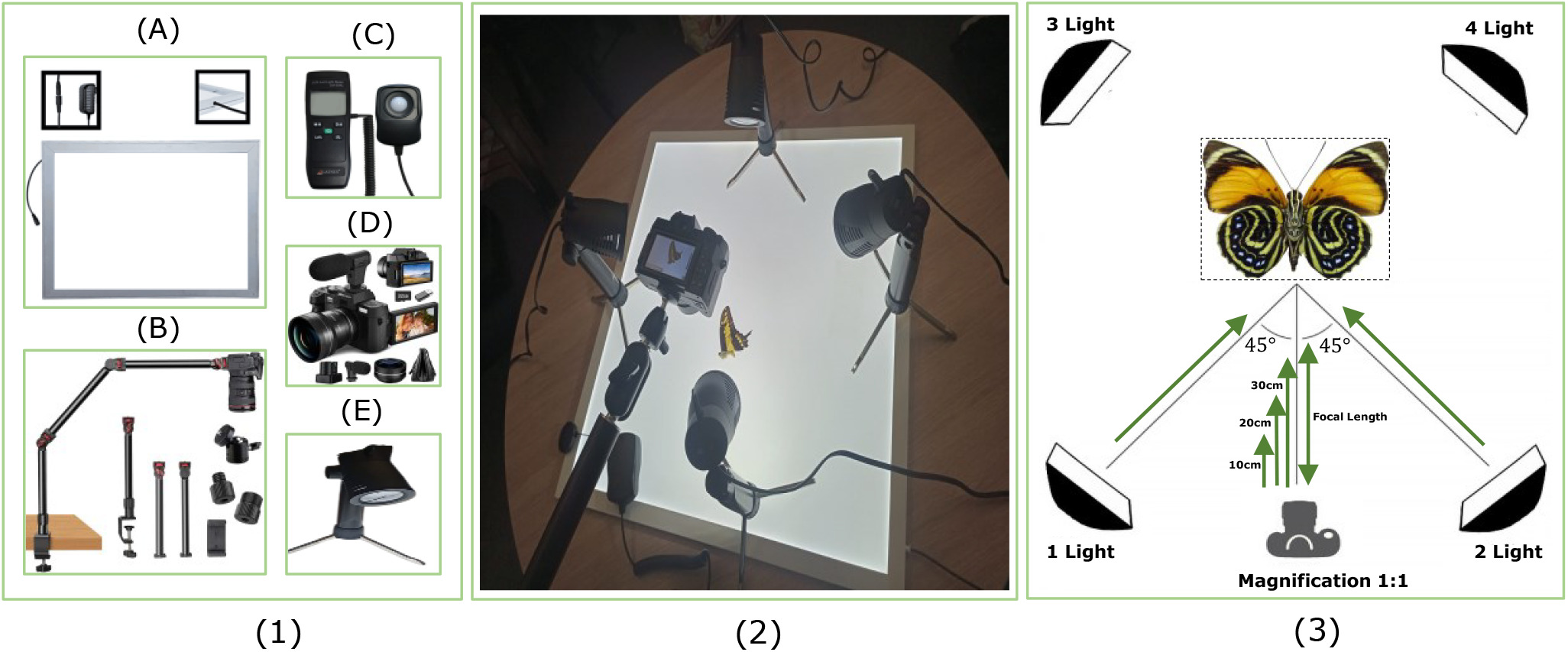
(1) Experimental Equipments: White Light Board (A), Tripod stand (B), LUX meter (C), DSLR camera Monitech4K UltraHD 48MP, featuring a 16x digital optical zoom lens with 52mm, f/5.04mm, wide-angle lens, (D), and 4 white light LED lamps (E). (2) Experimental setup, and (3) Experimental Parameters: The experiment was conducted using three different working distance lengths (10cm, 20cm, and 30cm) with 1:1 magnification to ensure consistent image capture and depth of field, and the lighting setup included four adjustable lamps positioned at 45° angles with the background and foreground lux of 100 to 650 with an interval of 50, while the camera was mounted at a 90° top-down orientation

We set up the camera parameters according to the macro lens specifications, using a 52mm lens at 1:1 magnification and an aperture of f/5.6, without applying any digital zoom, to ensure accurate and high-resolution imaging of fine morphological details. We furthermore chose three working distances at which to take images: 10cm, 20cm, and 30cm. The images were captured at a 1:1 magnification ratio and positioned at a 90° angle relative to the white light board. This angle was particularly useful for documenting the full body structure of insects, as it minimized distortion and ensured that both symmetry and fine morphological details, such as patterns on the wings or body segments, were visible. This wider aperture helped brighten the image, while the shallow depth of field enabled easy focus on specific insect features.

According to both museum standards and entomologists, 50 lux is a minimum requirement for very sensitive items such as insects, feathers, and fur, if color and detail are to be adequately preserved without compromise in visibility. However, some darker colors require illumination levels of up to 500 lux to ensure visible clarity. [60–63]. Some laboratory lighting guidelines for color inspection require illuminanation levels of 1000 lux to meet the visual accuracy standards [64, 65]. Given the vast variability in the available guidelines, we used varying light intensities ranging from 100 to 650 lux, as this range is representative for both background and foreground illumination.

We limited our experiment to a maximum of 650 lux, as results from our software (AInsectID Version 1.1 Color Merge) reveal that beyond 500 lux, the color difference-measured using the Δ*E*_00_ metric remains below 1, which is within the Just Noticeable Difference (JND) threshold. This indicates that a further increase in lux would have a negligible impact on perceptible color changes, which will be discussed in more detail in the Results section. Our experiment made use of four lamps to evaluate the effect of lighting conditions on the color merging algorithm. The first setup involved using a single light positioned at a 45^°^ angle, emitting light onto the specimen.

Images were captured at three working distances (10cm, 20cm, 30cm) from 100 to 650 lux. Specifically, the light was positioned at a 45^°^ angle to the top surface of the specimen. It was directed at a diagonal, creating an optimal angle of incidence as shown in Figure 2. This angle helped to evenly distribute the light across the specimen, minimizing dark shadows and reducing glare [66]. In the second setup, two lights were positioned on opposite sides of the specimen, each at a 45^°^ angle, and the same imaging procedure was repeated. For the third setup, the lights were adjusted to illuminate the specimen from three sides. Finally, in the fourth setup, four lights were used to ensure uniform illumination and cancel out any lighting effects from opposite directions. This systematic approach allowed us to assess the impact of varying light configurations on algorithm performance. These set-up tests also help us to capture the true color patterns of the specimen by preventing light from hitting the specimen directly from one angle. In the set-ups where there was more than one lamp used, the light crossed over from multiple directions, minimizing shadows, highlighting both the surface features and the color patterns without overwhelming the subject with glare or excessive reflection, and enhancing the visibility of color patterns.

- **Data Collection and Processing**

Five different butterfly wings were used to test the algorithm’s color-merging consistency, Figure 1. These wings were selected for their high representations of structural color, allowing us to assess how effectively the algorithm merges color patterns across varying wing types. For each insect wing, we processed different colors to assess the accuracy and consistency of the color merging algorithm as shown in Figure 3. These 16 colors were carefully selected to represent key aspects of each specimen’s color pattern. The AInsectID Version 1.1 Color Merge algorithm was applied to each image, and the resulting color data was analyzed to evaluate how well the algorithm handled color merging across varying lighting conditions and working distance. By processing multiple colors for each wing, we were able to gain a comprehensive understanding of the algorithm’s performance and its ability to capture and merge the color patterns of the specimen accurately. We captured five images of each wing for each setting, such as 100 lux and 10 cm working distance with one light lamp. These images were then processed using the AInsectID Version 1.1 Color Merge algorithm. After processing, we calculated the average of RGB values of each color across the five processed images, with this average taken as the final resulting RGB value. By averaging, we obtain a more reliable and representative color value for each setting. We prefer this approach over reliance on a single measurement, as it could be influenced by fluctuations.

**Figure 3.**
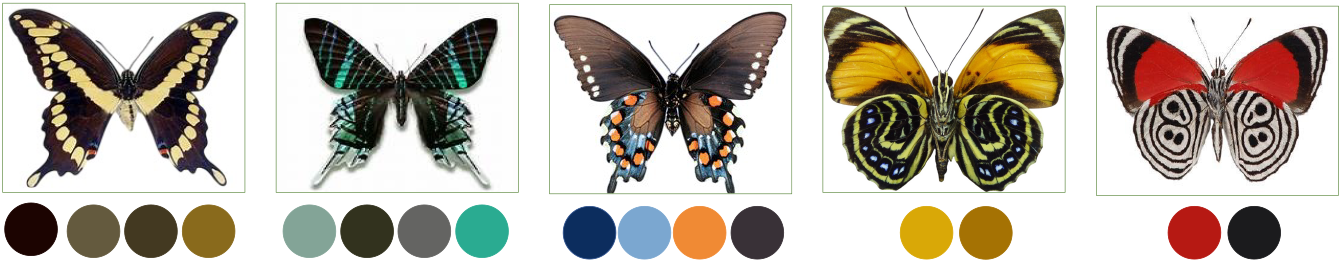
Color selection process: Five butterfly specimens were chosen, distinct colors are selected from each specimen for processing and analysis using the AInsectID Version 1.1 Color Merge algorithm

We repeated this process for each combination of lighting with working distance, to ensure consistency and accuracy could be achieved using the color merging algorithm. Specifically, 720 images were captured for each insect wing, using three working distances, four lighting positions, and light intensities ranging from 100 to 650 lux, with intervals of 50 lux. This procedure was repeated for five different insect wings, resulting in a total of 3600 images for testing and processing. As shown in Figure 4, we recorded 720 different RGB values for each color, independently varying lux intensities, number of lamps used, and working distances (expressed as focal lengths). With 16 colors, this resulted in a total of 11,520 RGB values.

**Figure 4.**
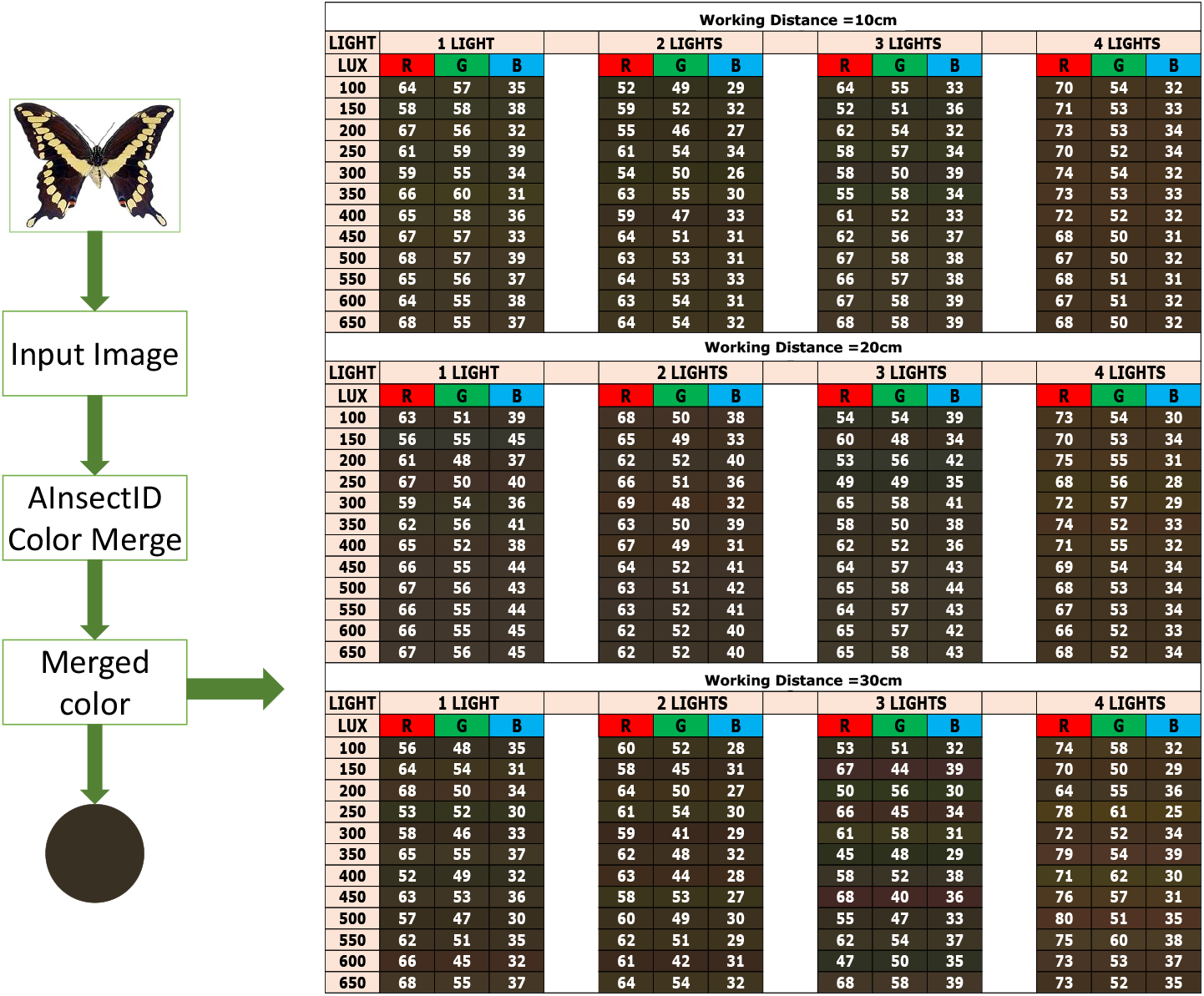
Color processing Workflow: In each experimental setup, five images were captured for each butterfly wing to ensure consistency (this approach is preferred over relying on a single measurement, as individual measurements could be influenced by fluctuations), resulting in a total of 720 (n = 720) images per wing and 3,600 images across five wings (n = 3600). All images were processed using the AInsectID Version 1.1 Color Merge algorithm to extract color information for 16 distinct colors. For each color, the RGB values from the five images (n = 5) were averaged to obtain a single representative RGB value. These averaged values were then used for subsequent analysis. In total, 720 distinct RGB values (n = 720) were recorded for each of the 16 colors, yielding 11,520 RGB measurements. Data collection was performed under varying conditions, including three working distances (10cm, 20cm, 30cm) with 1:1 magnification, four lighting positions, and light intensities ranging from 100 to 650 lux.

We using the Δ*E*_00_ function, as in Equation 1, with a 0.5 threshold. We analyze variations across lux levels (ranging from 100 to 650 in intervals of 50) for working distances of 10cm, 20cm, and 30cm. This is done separately for each setup, i.e. using 1 light, 2 lights, 3 lights, and 4 lights, to assess color consistency under different lighting conditions with a 1:1 magnification. Figure 5 illustrates the color difference with a single light source at a 10 cm working distance. Δ*E*_00_ less then 0.5 threshold allowed us to determine perceptible differences in color merging accuracy. By analyzing these differences, we identified the experimental setup that resulted in the least color variation, providing insights into the algorithm’s robustness and optimal imaging conditions for accurate color merging. After calculating the color differences for all setups, results are visualized using box and whisker plots and choropleth maps, with each plot representing the color variations for a single color under each lighting condition. This approach provided a comprehensive representation of variability across lighting conditions, helping to identify the experimental setups under which the color merging algorithm becomes less sensitive to lighting variations and produces the most consistent output.

**Figure 5.**
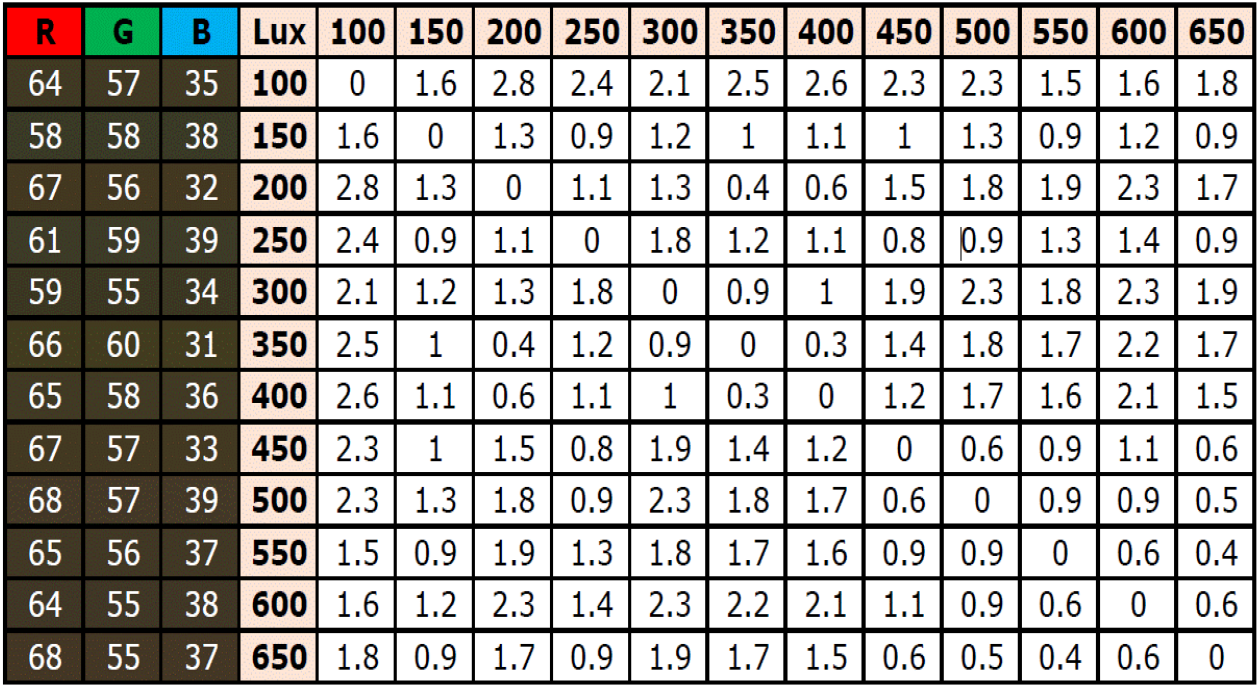
Illustration of the color difference calculation using the Δ*E*_00_ metric, applying a threshold of 0.5 to assess perceptual color variations. In each experimental setup, a total of 55 (n = 55) color difference values are computed, representing the pairwise Δ*E*_00_ values across each experimental setup. This results in a total of 660 (n = 660) Δ*E*_00_ values per color.

To determine whether the color merging algorithm maintains consistent color differences across various colors under different levels of illumination (100-650), we conducted a statistical analysis using the mean and standard deviation of the color difference metric (Δ*E*_00_). For each setup, we first computed the Δ*E*_00_ color differences individually for all 16 colors. We then calculated the mean and standard deviation (Δ*E*_00_) for all 16 colors cumulatively to assess both the average color merging performance and the variation across colors within that specific lighting condition and fixed working distance (10cm, 20cm, and 30cm) as shown in Equation 2 and 3 respectively. As we are working with 16 color samples denoted as *C*_1_, *C*_2_, …, *C*_16_, at each lux interval *L*_*j*_ ∈ {100–150, 150–200, …, 600–650}, we compute the color difference 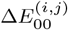 for each color *C*_*i*_, where *i* = 1, 2, …, 16 and *j* = 1, 2, …, 11. Then, at each lux interval *L*_*j*_, the mean and standard deviation of the color difference are calculated according to Equations 2 and 3, respectively.

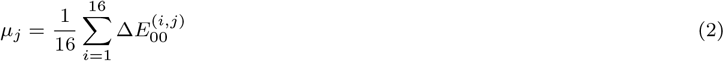

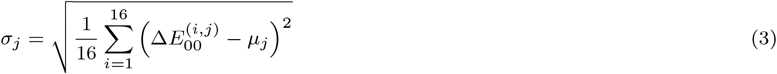

This approach allowed us to determine how consistently the algorithm handled different colors under each level of illumination, providing insight into its robustness and perceptual accuracy. Upon identifying an optimal setup, defined as the condition under which the mean Δ*E*_00_ ≤ 1 (Just Noticeable Difference, JND), we extended the evaluation to assess the robustness of the algorithm. This involved testing the system on more than 50 additional color samples under the same optimized conditions. To quantify the statistical significance and consistency of the results, we calculated confidence intervals at 90%, 95%, and 99% confidence levels. This allowed us to assess the reliability and stability of the algorithm’s color merging performance under real-world conditions, confirming that the software consistently maintained imperceptible color differences across a wider range of colors.

## 5 Results and Discussion

### 5.1 Performance Analysis of the Experimental Setup

This section presents and analyzes the outcomes of the study, focusing on the performance of the experimental setup and of the AInsectID Version 1.1 Color Merge algorithm under controlled lighting conditions. The findings are evaluated to assess the effectiveness of the color merging process, its consistency across different lighting intensities and working distances, and its alignment with established thresholds of human color perception. The results are presented using a series of box-and-whisker plots that illustrate the range of color differences (Δ*E*_00_) observed for each of the 16 colors under varying working distances (10 cm, 20 cm, and 30 cm) and lighting setups (1, 2, 3, and 4 lights), with corresponding illuminance levels ranging from 100 to 650 lux. The primary objective of this analysis is to identify the experimental setup that yields the lowest color differences range, indicating the most stable and consistent color merging. We have total 220 color difference (Δ*E*_00_) values (n = 200) for the experimental setup for each color.

Upon identifying the optimal setup, defined as the combination of working distance and lighting condition that consistently yields the lowest Δ*E*_00_ values, more detailed analyses and visualizations were conducted to gauge the influence of varying illumination levels (lux) on AInsectID’s color merging performance. This was visualized using a choropleth map, representing the distribution and intensity of color differences across different lux intervals, providing an intuitive understanding of the algorithm’s stability under varying lighting conditions. In the choropleth maps, each lux level is represented as a distinct region and is shaded according to its corresponding Δ*E*_00_ value. The purpose of exploring the data using a choropleth map is to identify the specific illuminance levels (lux) within the optimal experimental setup that result in color differences (Δ*E*_00_) below the JND threshold, defined as Δ*E*_00_ ≤ 1. At this threshold, color differences are imperceptible to the human eye and are therefore considered negligible, as previously discussed. This visualization facilitates a clear understanding of the optimal range of lux values within which color merging remains consistent and stable.

Figures 6, 7, 8, and 9 demonstrate that the 10 cm working distance consistently results in a narrower range of color differences as compared to the 20 cm and 30 cm working distances in analyses conducted on *Papilio glaucus, Philolaus, Battus philenor*, and *Agrias Amydon & Diaethris*, respectively. The setup using 4 light sources at a 10 cm distance shows the lowest and most stable Δ*E*_00_ values in every box and whisker plot. As indicated in the four box and whisker plots of Figure 6, the configuration with 4 lights at a 10 cm working distance consistently exhibits the minimum color difference ranges, specifically: 0.3231-2.8345, 0.6279-8.3810, 0.3640-6.0729, and 0.1238-2.4459, respectively. In contrast, the 30 cm working distance displays significantly higher variability and larger maximum color difference values. To further analyze this optimal setup (minimal color difference), a choropleth map was generated to visualize color variation (Δ*E*_00_) and its behavior across different lux intervals within this configuration, offering deeper insight into its performance under varying lighting conditions. In almost all the choropleth maps, the lower-right region consistently displays lower Δ*E*_00_ values, typically in the range 0 to 1.5 close to JND threshold (Δ*E*_00_ ≤ 1), which predominantly corresponds to the higher illumination levels ranging from 500 to 650 lux. This indicates that increased lighting intensity in this range contributes to more stable and minimal color differences during the merging process. similarly, in Figures 7, 8, and 9, the 10 cm working distance with 4-lights setup is most consistent in demonstrating the lowest color differences across the box and whisker plots. The corresponding choropleth maps for this optimal configuration predominantly display Δ*E*_00_ values close to or below JND threshold (Δ*E*_00_ ≤ 1), particularly concentrated within the 450–650 lux range. This finding indicates that when using lighting conditions within the 500 to 650 LUX range (background and foreground), the AInsectID Version 1.1 Color Merge algorithm performs optimally, producing results that are perceptually stable and in line with human visual sensitivity. Therefore, higher lux levels help to standardize color representation, making the output more reliable for color-based image analysis. We limited the maximum illumination level to 650 lux because our robustness testing, which will be discussed further in Section 5.3, indicated that at 550 lux, the color difference values (Δ*E*_00_) for all 16 colors consistently remained less than 1.0, which corresponds to the just noticeable difference (JND) threshold for perceptual color variations. To confirm stability, we further increased the illumination to 600 lux and 650 lux, and observed that the Δ*E*_00_ values continued to remain under 1.0 for all colors. This consistency indicates convergence in color merging performance, beyond which further increases in illumination did not yield significant differences. As such, 650 lux was chosen as the upper bound with a working distance of 10cm.

**Figure 6.**
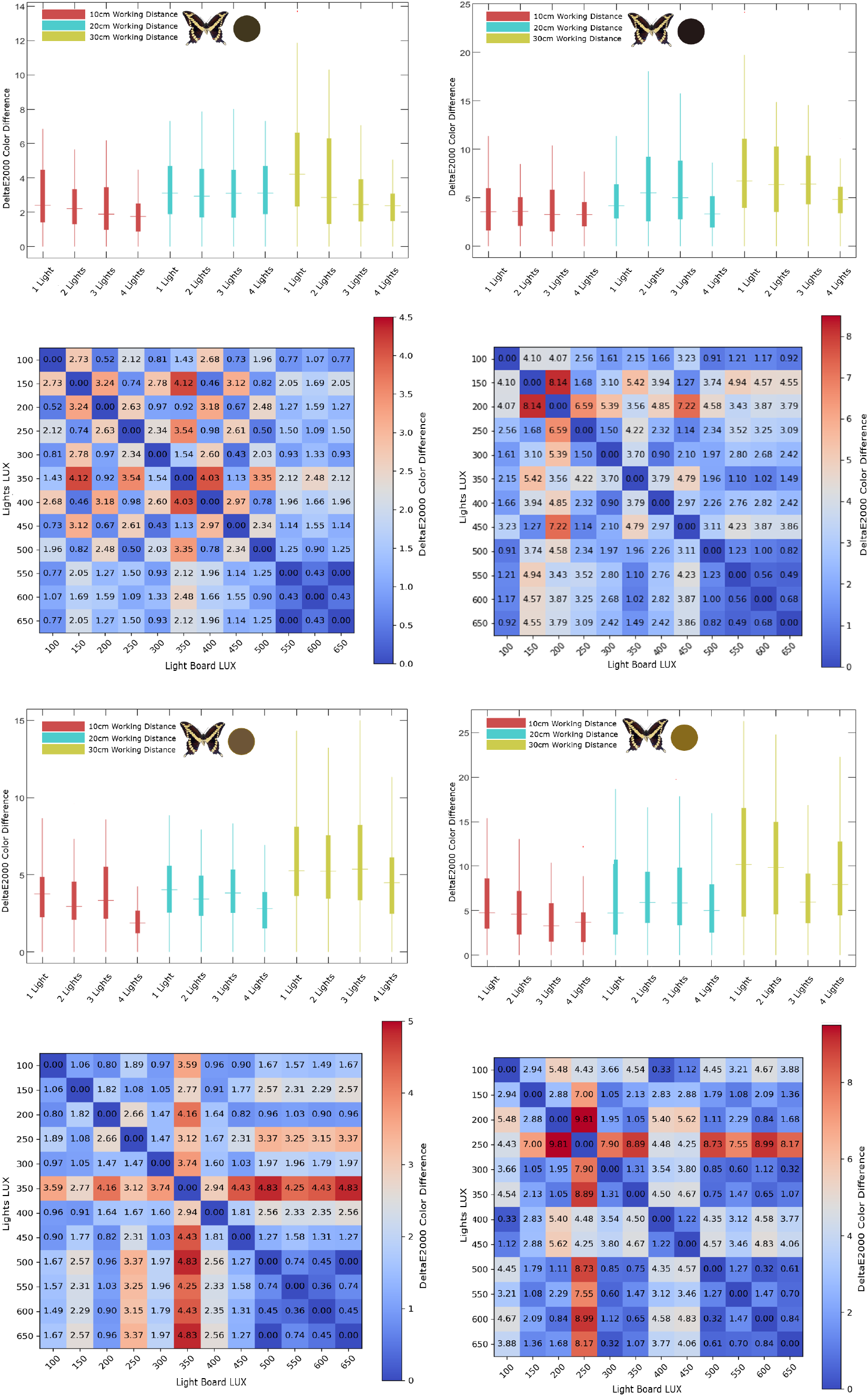
AInsectID Version 1.1 Color Merge was used to analyze color merging for *Papilio glaucus* under different lighting conditions (LUX) and working distances. For each setup, Δ*E*_00_ values (N=220) were obtained per wing. The box-and-whisker plots demonstrate that the 10 cm distance with 4 lights results in the smallest range of color differences. This finding is supported by choropleth maps, which identify specific lux levels associated with the most stable color performance

**Figure 7.**
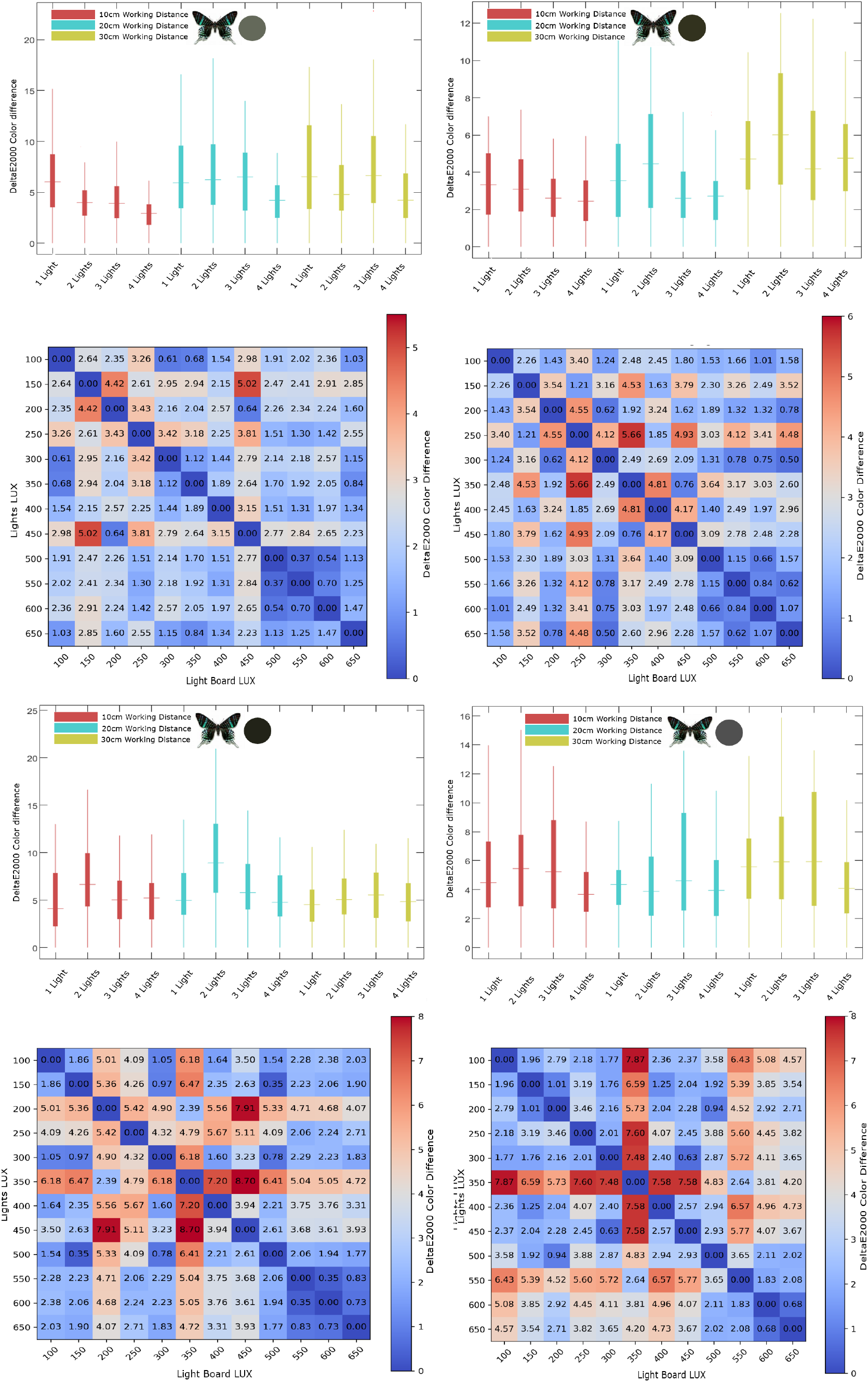
AInsectID Version 1.1 Color Merge was used to analyze color merging for *Philolaus* under different lighting conditions (LUX) and working distances. For each setup, Δ*E*_00_ values (N=220) were obtained per wing. The box-and-whisker plots demonstrate that the 10 cm distance with 4 lights results in the smallest range of color differences. This finding is supported by choropleth maps, which identify specific lux levels associated with the most stable color performance

**Figure 8.**
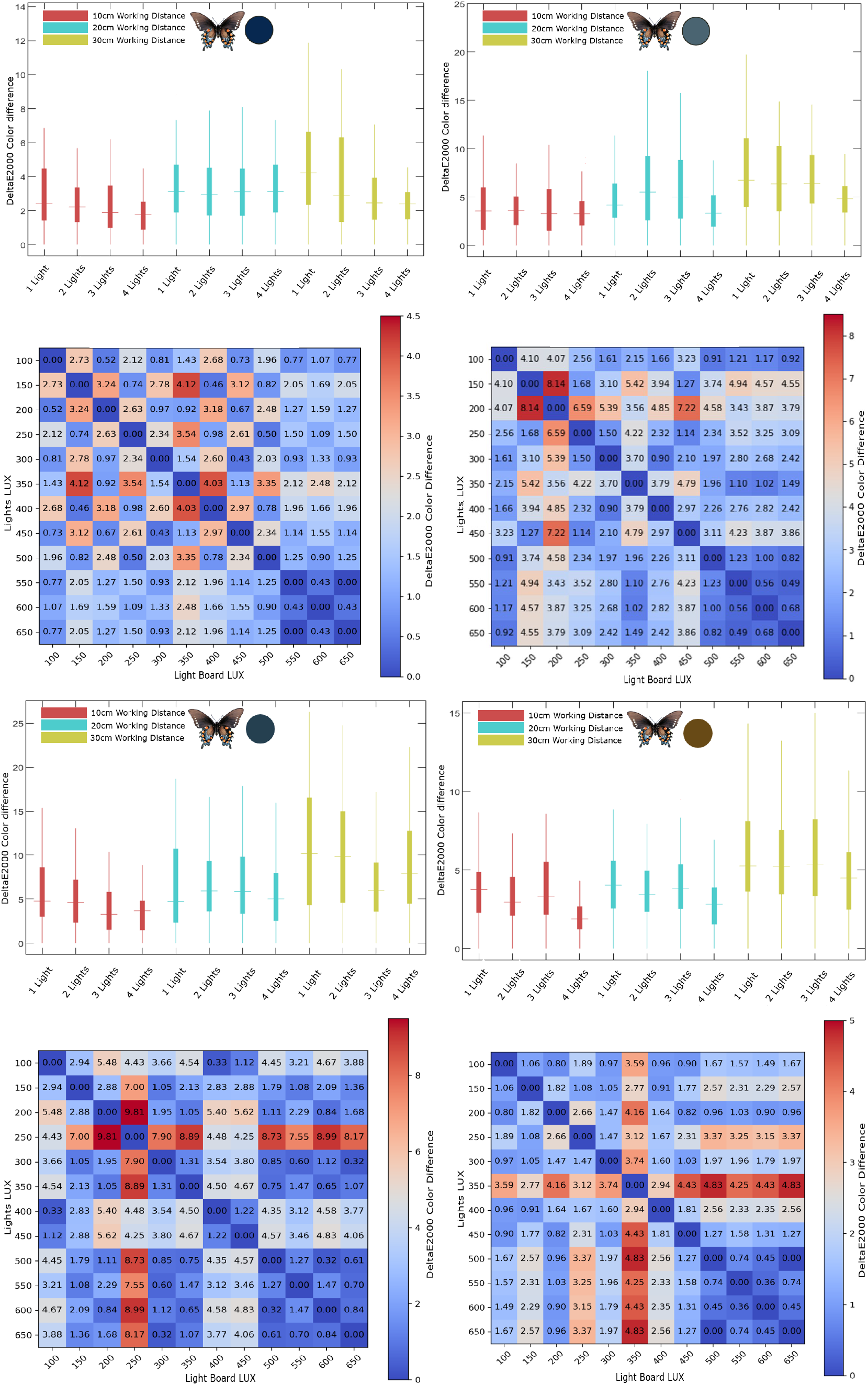
AInsectID Version 1.1 Color Merge was used to analyze color merging for *Battus philenor* under different lighting conditions (LUX) and working distances. For each setup, Δ*E*_00_ values (N=220) were obtained per wing. The box-and-whisker plots demonstrate that the 10 cm distance with 4 lights results in the smallest range13of color differences. This finding is supported by choropleth maps, which identify specific lux levels associated with the most stable color performance

**Figure 9.**
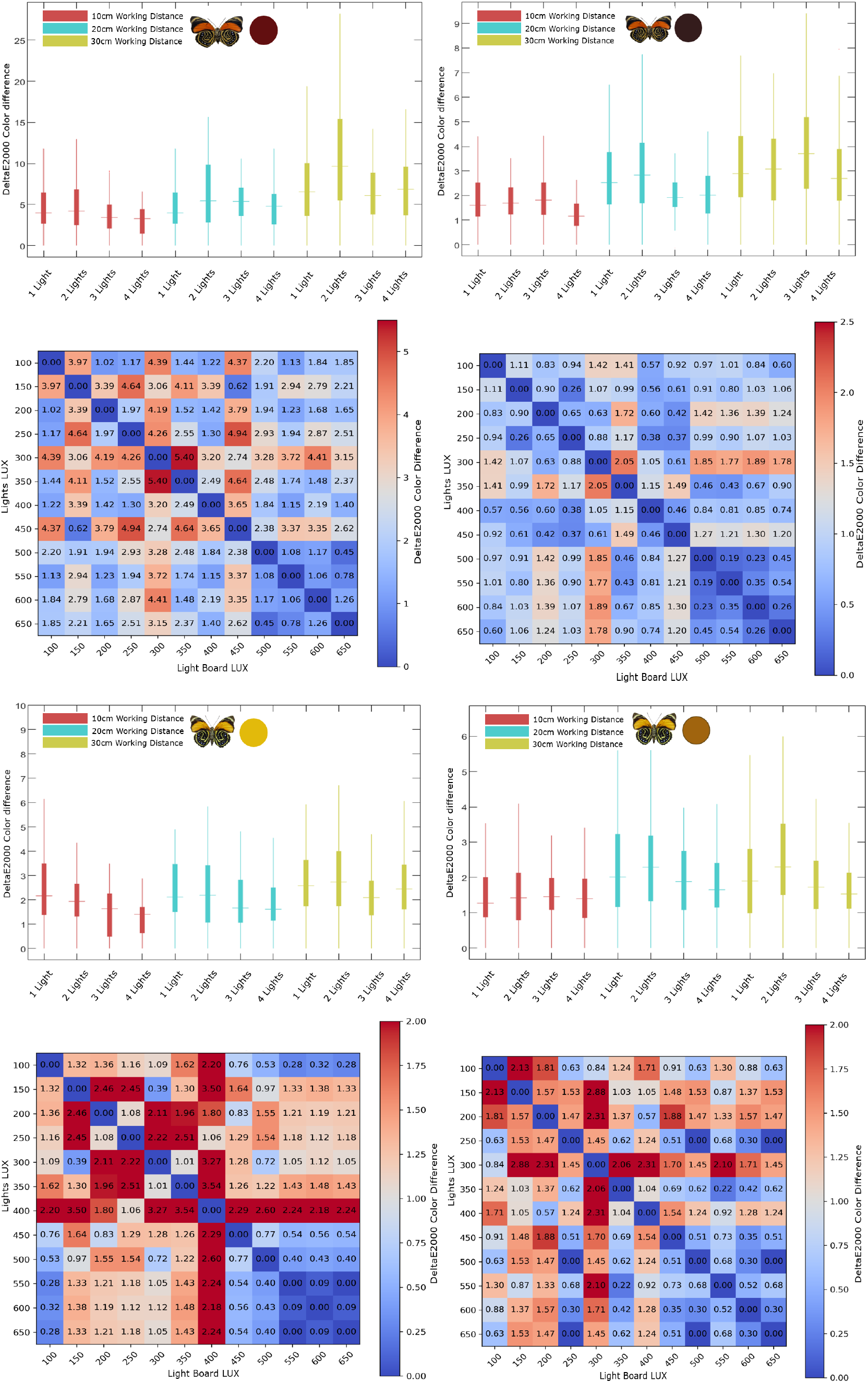
AInsectID Version 1.1 Color Merge was used to analyze color merging for *Agrias Amydon & Diaethris* under different lighting conditions (LUX) and working distances. For each setup, Δ*E*_00_ values (N=220) were obtained per wing. The box-and-whisker plots demonstrate that the 10 cm distance with 4 lights results in the smallest range of color differences. This finding is supported by choropleth maps, which identify specific lux levels associated with the most stable color performance

The analyses in Figures 7, 8, and 9 were conducted on an individual color and individual lux level basis. Tables 1, 2, and 3 provide further detail on the means and standard deviations of each color difference (Δ*E*_00_) across the full range of light intensities from 100 to 650 lux for each for each light source (Light 1, Light 2, Light 3, and Light 4), with a fixed working distance of 10cm, 20cm and 30cm respectively. A lower mean (Δ*E*_00_), particularly with values close to or below 1, indicates smaller perceptible average color differences, which implies more consistent color merging, while the standard deviation provides insight into the consistency of that performance across the different lux levels. As can be observed in Table 1, across the 16 colors, the setup of 4 lights at a fixed working distance of 10 cm consistently produces the lowest values of color difference, with means between 1.0706 and 1.9242, suggesting enhanced color accuracy and less perceptible deviation. In contrast, the 3-lights configuration often shows the highest mean (Δ*E*_00_) values, exceeding 4.2837 for several colors, indicating there are greater color deviations. The 1-light and 2-lights setups perform moderately, with mean values mostly in the 2.9043 to 3.6195 range. The standard deviations also tend to be lower in the 4-lights setup, indicating more consistent color rendering across all color samples. This suggests that increasing the number of lights not only improves the average color accuracy, but that it also stabilizes the color merging performance across different experimental conditions. Table 2 provides the mean and standard deviation values of color difference (Δ*E*_00_) for 16 colors at a 20 cm working distance. The 4-lights setup generally shows lower mean (Δ*E*_00_) values (2.5246 to 3.0229), indicating relatively enhanced color merging consistency. The 3-lights setup consistently has the highest color differences (4.9006 to 5.5787), while the 1-light and 2-lights setups performs moderately with ranges of (3.1656 to 4.0076) and (4.1979 to 4.9834), respectively, across all colors. Table 3, provides the mean and standard deviation of color differences (Δ*E*_00_) for 16 colors at a 30 cm working distance, shows that the 3-lights setup consistently yields the highest mean (Δ*E*_00_) values across most colors, indicating greater color inaccuracy. In contrast, the 4-lights setup yields lower (Δ*E*_00_) values (relatively), though still not as low as those observed at shorter working distances (e.g., 10 cm). The standard deviations are also generally higher at this distance, indicating greater variation in performance.

**Table 1:** Table shows mean ± standard deviation of Δ*E*_00_ values (n = 220 for each color under a 1, 2, 3 and 4-lighting setup) at a fixed working distance of 10 cm. A mean value close to 1 indicates high consistency in color merging, meaning the perceived color differences are minimal across varying lux (100-650)

| Working Distance = 10 cm |  |  |  |  |
| --- | --- | --- | --- | --- |
| Colors | 1 Light | 2 Lights | 3 Lights | 4 Lights |
| C1 | 3.6195± 1.6645 | 3.0004±0.8865 | 3.7338± 1.1524 | 1.7519± 0.1546 |
| C2 | 3.1384± 1.6798 | 2.9484± 0.8029 | 3.9393± 1.1940 | 1.6204± 0.1485 |
| C3 | 2.9555± 1.2864 | 3.1109± 0.1138 | 3.7789± 1.2456 | 1.7314± 0.0004 |
| C4 | 3.5549± 1.6821 | 3.1775± 0.1708 | 3.8319± 1.2621 | 1.7314± 0.0291 |
| C5 | 3.1934± 1.2548 | 3.0996± 0.9683 | 3.8807± 1.2393 | 1.8138± 0.0334 |
| C6 | 3.2524± 1.3791 | 3.1765± 0.8712 | 3.9447± 1.0829 | 1.8932± 0.2913 |
| C7 | 3.6508± 1.2350 | 3.1884± 0.9076 | 3.9998± 1.3372 | 1.5955± 0.1287 |
| C8 | 3.5383± 1.6076 | 3.0198± 0.1357 | 3.8174± 1.2117 | 1.0706± 0.3896 |
| C9 | 3.5208± 0.9366 | 2.8568± 0.7645 | 3.6965± 1.4702 | 1.3648± 0.2200 |
| C10 | 3.4880± 1.3359 | 3.0022± 0.8251 | 3.9752± 0.2439 | 1.6806± 0.1879 |
| C11 | 3.4845± 0.9920 | 3.0423± 0.0403 | 3.9150± 1.2498 | 1.6925± 0.2815 |
| C12 | 3.2824± 1.1886 | 3.3802± 0.0394 | 3.8159± 1.2793 | 1.6804± 0.0392 |
| C13 | 3.2824± 1.1108 | 3.1427± 0.6601 | 3.9134± 1.0582 | 1.7901± 0.2611 |
| C14 | 3.0115± 1.1528 | 3.1216± 0.9882 | 4.2837± 1.0552 | 1.7185± 0.0643 |
| C15 | 3.5834± 1.2252 | 3.2178± 0.8816 | 4.0616± 1.1051 | 1.9242± 0.2075 |
| C16 | 3.3719± 1.2442 | 2.9043± 0.7248 | 4.0889± 1.0570 | 1.8235± 0.0247 |

**Table 2:** Table shows mean ± standard deviation of Δ*E*_00_ values (n = 220 for each color under a 1, 2, 3 and 4-lighting setup) at a fixed working distance of 20 cm. A mean value close to 1 indicates high consistency in color merging, meaning the perceived color differences are minimal across varying lux (100-650)

| Working Distance = 20 cm |  |  |  |  |
| --- | --- | --- | --- | --- |
| Colors | 1 Light | 2 Lights | 3 Lights | 4 Lights |
| C1 | 4.5407±1.2379 | 4.0076±0.3408 | 5.2874±0.9691 | 2.5246±0.2638 |
| C2 | 4.5407±1.5954 | 3.7294±0.1859 | 5.5404±0.7022 | 2.9179±0.2592 |
| C3 | 4.2054±1.1328 | 3.6220±0.2939 | 4.6982±0.5860 | 2.8518±0.2441 |
| C4 | 4.3469±2.2513 | 3.3000±0.1160 | 5.4954±1.7898 | 2.6983±0.1989 |
| C5 | 4.9834± 1.7217 | 3.6211±1.3417 | 5.4972±0.79619 | 2.6962±0.1384 |
| C6 | 4.4673±1.5165 | 3.5566±0.3443 | 4.9250±0.8096 | 2.8207±0.1977 |
| C7 | 4.4490± 0.8303 | 3.9573±0.2950 | 4.9006±0.5805 | 3.1291±0.1490 |
| C8 | 4.1979±1.3934 | 3.5969±0.4005 | 5.2629±1.6855 | 2.5941±0.1675 |
| C9 | 4.4019±1.6715 | 3.6522±0.2744 | 5.3358±0.8641 | 2.8050±0.1414 |
| C10 | 4.3276±1.3874 | 3.3639±0.0281 | 5.5320±0.7577 | 2.8512±0.0188 |
| C11 | 4.5427±2.1803 | 3.5087±0.4044 | 5.3851±0.3018 | 2.7960±0.1472 |
| C12 | 4.3427±1.8761 | 3.4391±1.4622 | 5.0388±0.5695 | 3.0229±0.2531 |
| C13 | 4.4244±1.3329 | 3.4835±1.1481 | 5.3932±0.8043 | 2.7586±1.0034 |
| C14 | 4.5224±2.1351 | 3.1656±0.0625 | 5.2190±0.6516 | 2.9060±0.2727 |
| C15 | 4.6016±1.5826 | 3.3684±0.1832 | 5.5787±0.9978 | 2.7941±0.0294 |
| C16 | 4.7711±1.0880 | 3.4937±0.1254 | 5.2607±0.9548 | 2.6164±0.0294 |

**Table 3:** Table shows mean ± standard deviation of Δ*E*_00_ values (n = 220 for each color under a 1, 2, 3 and 4-lighting setup) at a fixed working distance of 30 cm. A mean value close to 1 indicates high consistency in color merging, meaning the perceived color differences are minimal across varying lux (100-650)

| Working Distance = 30 cm |  |  |  |  |
| --- | --- | --- | --- | --- |
| Colors | 1 Light | 2 Lights | 3 Lights | 4 Lights |
| C1 | 4.8112± 1.1232 | 4.6567± 1.3563 | 5.8268± 2.9572 | 3.3510±0.8261 |
| C2 | 4.8112± 2.2814 | 4.7997± 1.9125 | 5.9537± 2.5499 | 3.3310±1.1017 |
| C3 | 4.3290± 2.4053 | 4.8625± 1.8448 | 6.1829± 2.9955 | 3.4598±1.3576 |
| C4 | 4.1862±1.7596 | 4.9382± 1.6325 | 6.0154± 2.6779 | 3.3815±1.7804 |
| C5 | 4.1382± 1.1976 | 4.866± 1.4812 | 5.9033± 2.9506 | 3.3627±1.5465 |
| C6 | 4.0435± 1.9084 | 4.9713± 1.809 | 6.1842± 2.13248 | 3.4709±1.2782 |
| C7 | 5.0818± 2.3797 | 4.9810± 1.7054 | 5.7117± 2.6346 | 3.8063±1.2647 |
| C8 | 5.2147± 0.9527 | 4.7702± 1.6700 | 6.3072± 2.1472 | 3.6830±0.8576 |
| C9 | 4.1316± 1.4783 | 5.0729± 1.7342 | 5.9072± 1.1862 | 3.4938±1.1983 |
| C10 | 4.6216± 1.4246 | 5.1335± 1.6025 | 6.3429± 2.0062 | 3.5282±1.3838 |
| C11 | 4.4659± 2.3144 | 4.7828± 1.4622 | 6.3188± 2.7051 | 3.8951±1.6202 |
| C12 | 4.6259± 1.6472 | 5.1132± 1.6110 | 5.8491± 2.7345 | 3.7377±1.7343 |
| C13 | 4.9101± 2.4317 | 4.9909± 1.2573 | 5.94243± 2.1442 | 3.9926±1.5860 |
| C14 | 4.4867± 2.0140 | 4.9425± 1.4769 | 6.0538± 2.1523 | 3.8201±1.9476 |
| C15 | 4.8490± 1.5109 | 4.9891± 1.5886 | 5.8389± 2.1820 | 3.7257±1.3448 |
| C16 | 4.6243± 1.7547 | 4.7738± 1.4950 | 5.7145± 2.1980 | 3.6497±1.3273 |

Overall, the results indicate that the 4-light setup at a 10 cm working distance yields the most consistent color merging, yielding mean values between 1 and 1.9242. This suggests that images taken closer to the specimen contribute to more consistent color merging, likely due to the more controlled imaging environment minimizing optical distortions. The lower color variation at 10 cm can be attributed to enhanced precision in focus and reduced depth of field issues, which may have contributed to more accurate color representations.

Contrarily, the color variation increases significantly at a further distance (30cm). This variation was substantial enough that it often resulted in a shift to a different color (Δ*E* ≥ 6.0), highlighting the sensitivity of color perception to changes in working distance.

### 5.2 Robustness Testing of the Algorithm

To evaluate the robustness of the color merging performance of AInsectID Version 1.1 Color Merge under different variations in lux, we computed the mean and standard deviation of (Δ*E*_00_) color difference values cumulatively across all 16 representative color samples at a fixed working distance, Tables 4, 5 and 6. This analysis was performed to assess the overall performance of AInsectID Version 1.1 Color for all colors cumatively. Table 4 provides the mean ± standard deviation of (Δ*E*_00_) values measured across 16 color samples under four different lighting setups (1 light to 4 lights), with lux levels ranging from 100 to 650 lux at a fixed working distance of 10 cm. The results indicate substantial variation in color merging performance across both lighting conditions and lux ranges. At lower lux levels (100–450), 3 lights consistently exhibits the highest (Δ*E*_00_) values. For example, 5.8243 ± 0.4408 at 100–150 lux, suggesting less effective color merging. In contrast, 4 lights demonstrates superior performance, with (Δ*E*_00_) values progressively decreasing as illumination increases. By the time illumination reaches 450–500 lux, 4 lights achieves a mean color difference of 1.0419 ± 0.1071, approaching the perceptual threshold of 1.0. Most notably, from 500 lux onward, 4 lights consistently achieves (Δ*E*_00_) values below or equal to 1.0, with values of 0.7070 ± 0.0614, 0.5924 ± 0.0599, and 0.5927 ± 0.0500 across the 500–650 lux range. These results indicate that color differences under these conditions are imperceptible to the human eye, meeting the just noticeable difference (JND) threshold. Additionally, the low standard deviations observed under 4 lights at higher lux levels reflect highly consistent color merging across all 16 color samples. In contrast, 1 light through 3 lights continue to exhibit higher (Δ*E*_00_) values even at elevated illumination levels, suggesting less robust (Δ*E*_00_) values under 4 lights beyond 500 lux suggests convergence, indicating that further increases in lighting intensity do not yield noticeable improvements in perceptual color accuracy. These findings highlight 4 lights at 650 lux as the optimal condition for accurate and consistent color merging by AInsectID Version 1.1.

**Table 4:** Table showing mean ± standard deviation of color difference (Δ*E*_00_) measurements across 16 color samples under varying illumination conditions (100–650 lux) at fixed 10 cm working distance (n = 80 for each condition of lux with number of lights), indicating merging variation and consistency of AInsectID Version 1.1 color merging across 16 different colors. Mean (Δ*E*_00_) values below or equal to 1 indicate color differences are imperceptible to the human eye, demonstrating AInsectID’s high color merging accuracy. Low standard deviations reflect consistent performance across lighting conditions for different colors.

| Working Distance = 10 cm |  |  |  |  |
| --- | --- | --- | --- | --- |
| LUX | 1 Light | 2 Lights | 3 Lights | 4 Lights |
| 100–150 | 2.5270± 0.2971 | 4.1916±0.5562 | 5.8243± 0.4408 | 2.4645± 0.2330 |
| 150–200 | 2.3941± 0.8600 | 3.7778 ±0.52178 | 5.8149± 0.3020 | 3.3942± 0.2443 |
| 200–250 | 2.5496± 0.5338 | 3.3510± 0.5356 | 4.8866± 0.3292 | 2.8622± 0.2308 |
| 250–300 | 2.7336± 0.5954 | 3.4731± 0.3395 | 4.6120± 0.4274 | 2.8587± 0.4117 |
| 300–350 | 1.9336± 0.6515 | 3.1534± 0.4443 | 4.6120± 0.4274 | 1.8711± 0.2684 |
| 350–400 | 2.5401± 0.5303 | 3.1261± 0.3872 | 3.6189± 0.4768 | 1.7277± 0.1937 |
| 400–450 | 1.5689± 0.4866 | 2.9161± 0.2927 | 3.6189± 0.4768 | 1.4439± 0.1917 |
| 450–500 | 1.1544± 0.4202 | 3.0217± 0.3329 | 3.8332± 0.4249 | 1.0419± 0.1071 |
| 500–550 | 3.5875±0.55478 | 3.9949± 0.5964 | 3.7849± 0.33603 | **0.7070± 0.0614** |
| 550–600 | 3.5875±0.55478 | 2.4561± 0.2901 | 2.6288± 0.4351 | **0.5924± 0.0599** |
| 600–650 | 2.1612 ±0.72496 | 2.4309± 0.2928 | 2.4805± 0.4144 | **0.5927± 0.0500** |

**Table 5:** Table showing mean ± standard deviation of color difference (Δ*E*_00_) measurements across 16 color samples under varying illumination conditions (100–650 lux) at fixed 20 cm working distance (n = 80 for each condition of lux with number of lights), indicating merging variation and consistency of AInsectID Version 1.1 color merging across 16 different colors. Mean (Δ*E*_00_) values below or equal to 1 indicate color differences are imperceptible to the human eye, demonstrating AInsectID’s high color merging accuracy. Low standard deviations reflect consistent performance across lighting conditions for different colors.

| Working Distance = 20 cm |  |  |  |  |
| --- | --- | --- | --- | --- |
| LUX | 1 Light | 2 Lights | 3 Lights | 4 Lights |
| 100–150 | 6.2809±1.4246 | 5.3628±1.2929 | 6.4725± 0.6998 | 4.9638± 0.5959 |
| 150–200 | 6.6469±1.2221 | 5.9418±1.3119 | 6.0365± 0.6582 | 4.9591± 0.5459 |
| 200–250 | 6.2447±1.8012 | 4.5814±0.8232 | 5.2165± 0.5942 | 4.0246± 0.4588 |
| 250–300 | 4.0135±1.0045 | 4.5960±0.8230 | 5.03576± 0.6650 | 4.0345± 0.4485 |
| 300–350 | 4.4804±0.9227 | 4.1196±0.6778 | 5.3669± 0.7030 | 4.0523± 0.3125 |
| 350–400 | 4.3865±1.297 | 4.0949±0.7983 | 5.0447± 0.7244 | 3.9255± 0.3033 |
| 400–450 | 3.6008±0.5938 | 3.9648±0.5808 | 5.2366± 0.7211 | 3.9079± 0.3001 |
| 450–500 | 3.2291±0.8956 | 3.5232± 0.5346 | 4.3702± 0.7054 | 2.1351± 0.2512 |
| 500–550 | 3.5479±0.6512 | 3.3736± 0.4445 | 3.7778± 0.4214 | 2.0844± 0.2430 |
| 550–600 | 3.172±0.5815 | 3.2529±0.4272 | 3.7845± 0.4743 | 1.7251± 0.2411 |
| 600–650 | 3.6848±0.7587 | 2.6744±0.3959 | 3.6490± 0.4725 | **1.0009± 0.2080** |

**Table 6:** Table showing mean ± standard deviation of color difference (Δ*E*_00_) measurements across 16 color samples under varying illumination conditions (100–650 lux) at fixed 30 cm working distance (n = 80 for each condition of lux with number of lights), indicating merging variation and consistency of AInsectID Version 1.1 color merging across 16 different colors. Mean (Δ*E*_00_) values below or equal to 1 indicate color differences are imperceptible to the human eye, demonstrating AInsectID’s high color merging accuracy. Low standard deviations reflect consistent performance across lighting conditions for different colors.

| Working Distance = 30 cm |  |  |  |  |
| --- | --- | --- | --- | --- |
| LUX | 1 Light | 2 Lights | 3 Lights | 4 Lights |
| 100–150 | 7.2705±2.1528 | 6.8298± 1.5877 | 6.9912± 1.9264 | 5.4789± 0.6865 |
| 150–200 | 7.1877±2.1514 | 6.7881± 1.8251 | 6.0275± 0.6023 | 5.9773± 1.1352 |
| 200–250 | 7.2075±2.1041 | 6.1525± 1.4741 | 5.4177± 0.7437 | 5.645± 0.6649 |
| 250–300 | 7.5894±1.9816 | 6.9231± 1.3962 | 5.1585± 0.5972 | 5.4644± 0.3577 |
| 300–350 | 6.1185±1.2097 | 6.4172± 1.3307 | 5.3763± 0.6642 | 4.1184± 0.1712 |
| 350–400 | 6.0978±1.6737 | 5.2279± 0.4027 | 5.2707± 0.6758 | 4.0977± 0.6205 |
| 400–450 | 5.6673±1.4293 | 4.5900± 0.4682 | 4.3046± 0.6989 | 3.6672± 0.6838 |
| 450–500 | 4.9684±1.1372 | 4.0078± 0.4465 | 4.3346± 0.7356 | 3.5283± 0.7841 |
| 500–550 | 4.9032±1.9872 | 3.3263± 0.3588 | 3.1730± 0.7740 | 2.3200± 0.6797 |
| 550–600 | 3.8732±1.2132 | 3.19982± 0.3496 | 3.4808± 0.6855 | 2.8362± 0.32843 |
| 600–650 | 3.1542±1.7144 | 3.0655± 0.4575 | 3.5012± 0.7366 | 2.3394± 0.4290 |

Table 5 shows the mean ± standard deviation of color difference (Δ*E*_00_) at a working distance of 20 cm. Compared to the values shown in Table 4, which are at 10 cm working distance, the results in this table show generally higher color difference values, indicating less effective color merging performance at this longer working distance. At lower lux levels (100–400), all lighting configurations produce Δ*E*_00_ values significantly greater than the perceptual threshold of 1, with 3 lights often performing the worst (e.g., 6.4725 ± 0.6998 at 100–150 lux). 4 lights consistently achieves the lowest Δ*E*_00_ values among all setups, especially as lux increases. At 600–650 lux, only 4 lights approaches the perceptual limit with a value of 1.0009 ± 0.2080, showing a clear convergence trend similar to the 10 cm setup, though slightly less robust.

Table 6 provides the mean ± standard deviation values of color difference (Δ*E*_00_) across 16 color samples at a fixed working distance of 30 cm but at varying lux. When compared to the previous tables 4 and 5 (10 cm and 20 cm working distances, respectively), this setup shows a clear decline in color merging performance, as indicated by consistently higher Δ*E*_00_ values across all lighting configurations.

At the lower illumination range (100–300 lux), color difference values are notably high across all lighting setups, often exceeding 6 or 7 units for configurations with 1–3 lights. Even with 4 lights, which previously performed well, records values above 5 at the lower lux levels, far beyond the perceptual threshold of Δ*E*_00_ = 1. As illumination increases (400–650 lux), the color differences gradually decrease, particularly under the 4 lights configuration. However, even at the highest lux level (600–650), 4 lights only achieves a mean of 2.3394 ± 0.4290, which still does not reach the just noticeable difference (JND) threshold, unlike the 10 cm and 20 cm setups, where 4 lights converges to imperceptible color difference levels by 650 lux. This comparison of robustness highlights the importance of working distance, with the 10 cm condition enabling imperceptible color differences (Δ*E*_00_ ≤ 1) more reliably and at lower lux levels (starting from 500 lux), while the 20 cm and 30cm distances barely meet the JND threshold.

### 5.3 Critical Evaluation of Results

Based on our experimental findings, we identified that the optimal lux range for minimizing color differences in AInsectID Version 1.1 Color Merge falls between 500 and 650 lux. At these illumination levels, the color difference (Δ*E*) remains below 1 for certain colors, indicating a minimal perceptual difference between merged colors. To further evaluate the reliability and consistency of these findings, we conducted an extended analysis by testing an additional 50 colors within the identified lux range. This step was a crucial step to ensuring that the observed effect was not limited to a specific subset of colors but rather applied broadly across diverse color samples. The results, summarized in Table 7, present color samples with their RGB values and corresponding Δ*E* values at different lux ranges (500–550, 500–600, 500–650, 550–600, 550–650, and 600–650). The data indicates that the majority of colors processed by the merge algorithm exhibit a Δ*E* below 1 within specific lux conditions, highlighting the effectiveness of the 550–650 lux range in minimizing perceptual color differences. This suggests that under these optimal lighting conditions, the algorithm consistently achieves accurate color merging with minimal noticeable variation. Many colors maintain consistently low Δ*E* values across multiple lux intervals, suggesting stable color representation under these lighting conditions. For example, several colors, such as (153,113,86), (117,95,77), and (31,27,19), consistently show Δ*E* values below or equal to 1 in key lux ranges, reinforcing the suitability of this illumination range for accurate color merging. While most colors adhere to this trend, a few, such as (95,121,126) and (75,52,41), exhibit slightly higher Δ*E* values under broader lux variations, indicating that certain colors may be more sensitive to lighting changes. This suggests that 500–650 lux is generally optimal. Additionally, we calculate the level of confidence for our results to ensure statistical reliability of color merging algorithm and assess the significance of the observed color differences under different lux conditions. The measurements are evaluated under three distinct confidence levels: 90%, 95%, and 99%. These confidence levels indicate the certainty that the color differences fall within the observed ranges across repeated experiments. Table 8 demonstrates the color differences (Δ*E*) at various lux levels, with the goal of ensuring that these differences remain within the JND range, where Δ*E* less then and equal to 1. The results show that for various lux levels tested, particularly those between 550-650 lux, the Δ*E* values consistently fall below the JND threshold. At the 90% confidence level, color differences at lux levels such as 550-600, 550-650, and 600-650 are all within the range of 0.5 to 0.9 Δ*E*, which is well below the perceptible threshold of 1. These values are even more consistent at higher confidence levels (95% and 99%), further confirming that the observed color differences are unlikely to be noticeable by the human eye. For example, at the 550-600 lux level, the Δ*E* values at the 90%, 95%, and 99% confidence levels are between 0.6988 and 0.8596, all of which are well within the JND range. In contrast, at the 500-600 lux and 500-550 lux levels, the color differences show slightly higher Δ*E* values, but even at the 99% confidence level, these values remain just under the threshold of 1, indicating that the differences are still marginally perceptible or negligible in most practical scenarios. These results suggest that, under the specified lux levels, the color differences are statistically consistent and imperceptible to the human eye, validating the effectiveness of these lux levels in maintaining color merging algorithm consistency with minimal perceptual variation.

**Table 7:**
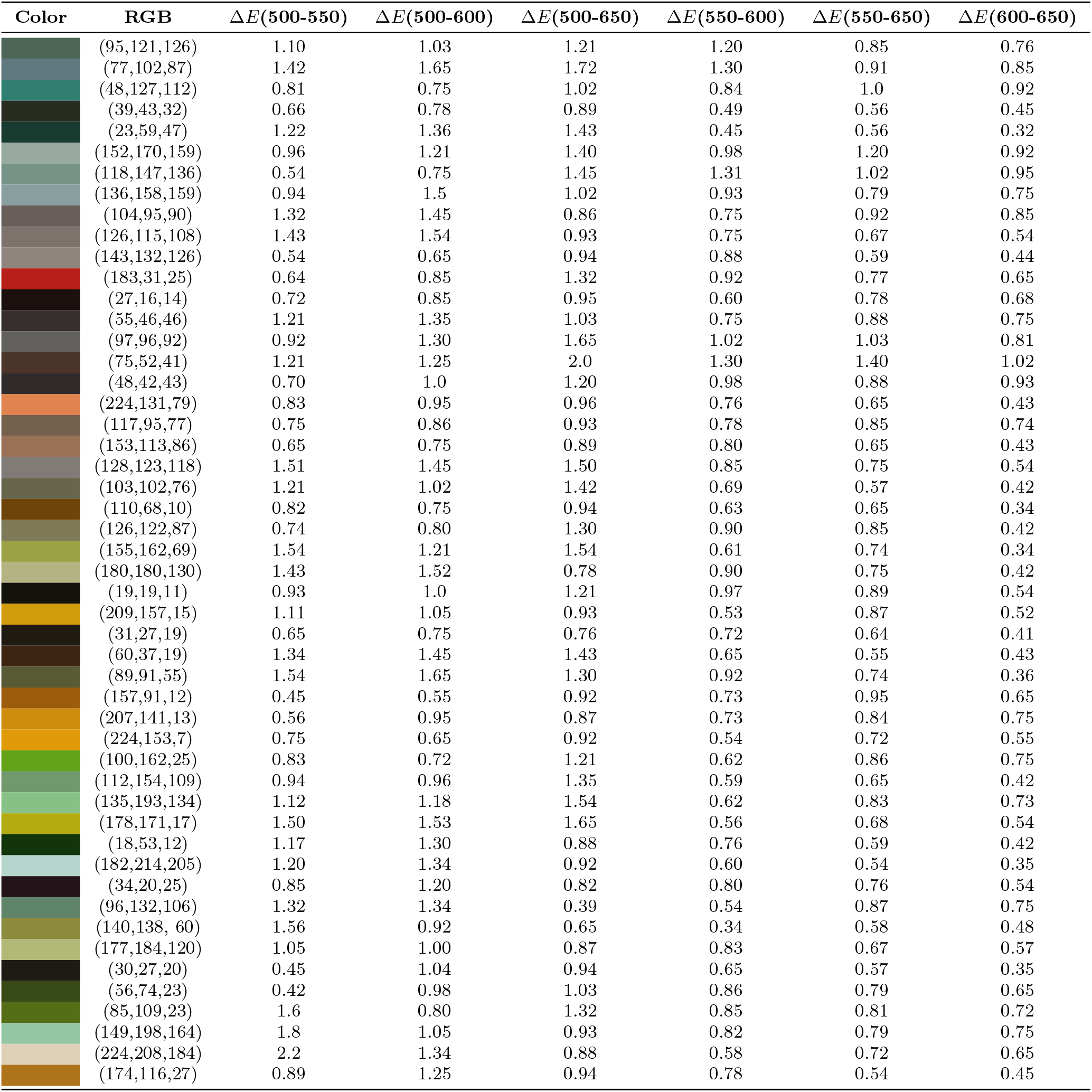
Color Differences Measured at Optimal Lux Levels Using Δ*E*2000: This table shows color differences (Δ*E*) at optimal Lux levels (500-550, 500-600, 500-650, 550-600, 550-650, and 600-650). For these Lux values, AInsectID Version 1.1 Color Merge algorithm shows color differences of less than 1, meeting the Just Noticeable Difference (JND) criteria. The Lux values are validated across 50 additional color samples, ensuring reliable and consistent results for color assessments in various lighting conditions

| Color | RGB | $\Delta E(500-550)$ | $\Delta E(500-600)$ | $\Delta E(500-650)$ | $\Delta E(550-600)$ | $\Delta E(550-650)$ | $\Delta E(600-650)$ |
| --- | --- | --- | --- | --- | --- | --- | --- |
|  | (95,121,126) | 1.10 | 1.03 | 1.21 | 1.20 | 0.85 | 0.76 |
|  | (77,102,87) | 1.42 | 1.65 | 1.72 | 1.30 | 0.91 | 0.85 |
|  | (48,127,112) | 0.81 | 0.75 | 1.02 | 0.84 | 1.0 | 0.92 |
|  | (39,43,32) | 0.66 | 0.78 | 0.89 | 0.49 | 0.56 | 0.45 |
|  | (23,59,47) | 1.22 | 1.36 | 1.43 | 0.45 | 0.56 | 0.32 |
|  | (152,170,159) | 0.96 | 1.21 | 1.40 | 0.98 | 1.20 | 0.92 |
|  | (118,147,136) | 0.54 | 0.75 | 1.45 | 1.31 | 1.02 | 0.95 |
|  | (136,158,159) | 0.94 | 1.5 | 1.02 | 0.93 | 0.79 | 0.75 |
|  | (104,95,90) | 1.32 | 1.45 | 0.86 | 0.75 | 0.92 | 0.85 |
|  | (126,115,108) | 1.43 | 1.54 | 0.93 | 0.75 | 0.67 | 0.54 |
|  | (143,132,126) | 0.54 | 0.65 | 0.94 | 0.88 | 0.59 | 0.44 |
|  | (183,31,25) | 0.64 | 0.85 | 1.32 | 0.92 | 0.77 | 0.65 |
|  | (27,16,14) | 0.72 | 0.85 | 0.95 | 0.60 | 0.78 | 0.68 |
|  | (55,46,46) | 1.21 | 1.35 | 1.03 | 0.75 | 0.88 | 0.75 |
|  | (97,96,92) | 0.92 | 1.30 | 1.65 | 1.02 | 1.03 | 0.81 |
|  | (75,52,41) | 1.21 | 1.25 | 2.0 | 1.30 | 1.40 | 1.02 |
|  | (48,42,43) | 0.70 | 1.0 | 1.20 | 0.98 | 0.88 | 0.93 |
|  | (224,131,79) | 0.83 | 0.95 | 0.96 | 0.76 | 0.65 | 0.43 |
|  | (117,95,77) | 0.75 | 0.86 | 0.93 | 0.78 | 0.85 | 0.74 |
|  | (153,113,86) | 0.65 | 0.75 | 0.89 | 0.80 | 0.65 | 0.43 |
|  | (128,123,118) | 1.51 | 1.45 | 1.50 | 0.85 | 0.75 | 0.54 |
|  | (103,102,76) | 1.21 | 1.02 | 1.42 | 0.69 | 0.57 | 0.42 |
|  | (110,68,10) | 0.82 | 0.75 | 0.94 | 0.63 | 0.65 | 0.34 |
|  | (126,122,87) | 0.74 | 0.80 | 1.30 | 0.90 | 0.85 | 0.42 |
|  | (155,162,69) | 1.54 | 1.21 | 1.54 | 0.61 | 0.74 | 0.34 |
|  | (180,180,130) | 1.43 | 1.52 | 0.78 | 0.90 | 0.75 | 0.42 |
|  | (19,19,11) | 0.93 | 1.0 | 1.21 | 0.97 | 0.89 | 0.54 |
|  | (209,157,15) | 1.11 | 1.05 | 0.93 | 0.53 | 0.87 | 0.52 |
|  | (31,27,19) | 0.65 | 0.75 | 0.76 | 0.72 | 0.64 | 0.41 |
|  | (60,37,19) | 1.34 | 1.45 | 1.43 | 0.65 | 0.55 | 0.43 |
|  | (89,91,55) | 1.54 | 1.65 | 1.30 | 0.92 | 0.74 | 0.36 |
|  | (157,91,12) | 0.45 | 0.55 | 0.92 | 0.73 | 0.95 | 0.65 |
|  | (207,141,13) | 0.56 | 0.95 | 0.87 | 0.73 | 0.84 | 0.75 |
|  | (224,153,7) | 0.75 | 0.65 | 0.92 | 0.54 | 0.72 | 0.55 |
|  | (100,162,25) | 0.83 | 0.72 | 1.21 | 0.62 | 0.86 | 0.75 |
|  | (112,154,109) | 0.94 | 0.96 | 1.35 | 0.59 | 0.65 | 0.42 |
|  | (135,193,134) | 1.12 | 1.18 | 1.54 | 0.62 | 0.83 | 0.73 |
|  | (178,171,17) | 1.50 | 1.53 | 1.65 | 0.56 | 0.68 | 0.54 |
|  | (18,53,12) | 1.17 | 1.30 | 0.88 | 0.76 | 0.59 | 0.42 |
|  | (182,214,205) | 1.20 | 1.34 | 0.92 | 0.60 | 0.54 | 0.35 |
|  | (34,20,25) | 0.85 | 1.20 | 0.82 | 0.80 | 0.76 | 0.54 |
|  | (96,132,106) | 1.32 | 1.34 | 0.39 | 0.54 | 0.87 | 0.75 |
|  | (140,138, 60) | 1.56 | 0.92 | 0.65 | 0.34 | 0.58 | 0.48 |
|  | (177,184,120) | 1.05 | 1.00 | 0.87 | 0.83 | 0.67 | 0.57 |
|  | (30,27,20) | 0.45 | 1.04 | 0.94 | 0.65 | 0.57 | 0.35 |
|  | (56,74,23) | 0.42 | 0.98 | 1.03 | 0.86 | 0.79 | 0.65 |
|  | (85,109,23) | 1.6 | 0.80 | 1.32 | 0.85 | 0.81 | 0.72 |
|  | (149,198,164) | 1.8 | 1.05 | 0.93 | 0.82 | 0.79 | 0.75 |
|  | (224,208,184) | 2.2 | 1.34 | 0.88 | 0.58 | 0.72 | 0.65 |
|  | (174,116,27) | 0.89 | 1.25 | 0.94 | 0.78 | 0.54 | 0.45 |

**Table 8:** Confidence level of Color Difference (Δ*E*) less then equal to 1: This table demonstrates color differences (Δ*E*) at optimal Lux levels, with a threshold of 1 to ensure that the differences remain within the Just Noticeable Difference (JND) range. The confidence levels for these measurements are set at 90%, 95%, and 99%, providing degrees of certainty in the consistency of color perception under optimal LUX levels

| Lux levels | 90% Significance level | 95% Significance level | 99% Significance level |
| --- | --- | --- | --- |
| 500-550 | 0.9502, 1.1370 | 0.9317, 1.1555 | 0.8944, 1.1928 |
| 500-600 | 1.0218, 1.1614 | 1.0080, 1.1752 | 0.9801, 1.2031 |
| 500-650 | 1.0440, 1.1936 | 1.0291, 1.2085 | 0.9992, 1.2384 |
| 550-600 | **0.6988, 0.8596** | **0.7189, 0.8395** | **0.7289, 0.8295** |
| 550-650 | **0.7333, 0.8155** | **0.7252, 0.8236** | **0.7088, 0.8400** |
| 600-650 | **0.5559, 0.6513** | **0.5464, 0.6608** | **0.5273, 0.6799** |

### 5.4 Future Considerations

While this study established an optimal setup for minimizing color variation in insect photography under controlled conditions, there are several avenues for future research. One key direction is expanding the evaluation to include a wider range of colors and insect species to assess an ability to generalize the AInsectID Version 1.1 Color Merge algorithm. Additionally, testing the algorithm with different camera models, image resolutions, and sensor types would provide insights into how hardware variations influence color accuracy. Another important extension is in transitioning from a controlled environment to real-world, open-light conditions. By evaluating the impact of natural and variable lighting scenarios, we can refine the algorithm to adapt to diverse environmental settings, improving its robustness and applicability in fieldwork. Furthermore, integrating advanced machine learning techniques could enhance the model’s ability to compensate for lighting inconsistencies, further improving color-based analyses. These future advancements will contribute to developing a more versatile and reliable system for insect photography and image processing.

## 6 Conclusions

This study evaluated the impact of various lighting conditions and working distances in a controlled environment on the performance of the AInsectID Version 1.1 Color Merge algorithm. Our findings highlight the optimal imaging setup required to minimize color variation in insect photography and this is illustrated in Appendix A1. Specifically, the optimal results were achieved using four white LED lights, positioned in a four corner arrangement (at 90° to one another) and further positioned at 45° angles relative to the specimen plane. This is to achieve balanced illumination with surface lux levels of 550–650 and a light board with a white background, maintaining an illumination range of 550–650 lux. This configuration ensures balanced illumination by allowing light sources to cancel out directional shadows. The camera is mounted directly above the specimen enabling a plan-view capture of the specimen perpendicular to the imaging surface, equipped with a 52mm macro lens, f/5.6 aperture, and 1:1 magnification at a 10 cm working distance, with ambient room lighting set to zero. While a plan view is used, the camera should remain fixed in this position relative to the specimen surface, with no tilt. However, it may rotate 360 ° around its vertical axis to adjust orientation, while maintaining a consistent perpendicular viewing angle. Capture resolution of the original image should be at least 2048×2048 px (4 MP). This configuration consistently yielded the lowest color variation, ensuring superior color consistency in AInsectID Version 1.1 Color Merge analysis. This lighting setup effectively neutralized external light interference, providing stable and uniform illumination, which is essential in obtaining reliable results. Our analysis of different lux levels confirmed that intensities within the 550–650 lux range produced color variations within the Just Noticeable Difference (JND) threshold, indicating perceptually stable outcomes. The imaging was performed without digital zoom, using a 1:1 magnification ratio to ensure accurate, true-to-scale top-down capture of the specimen. By adhering to this optimized setup, AInsectID Version 1.1 Color Merge ensures consistent and accurate color representation, enhancing the reliability of its color merging performance. This study provides a valuable framework for future research in insect photography and the development of advanced color-based image processing algorithms. While this algorithm performs optimally under controlled conditions, its performance may be uncertain if the recommended setup is not followed. Deviations from this optimal configuration may introduce variability in the results. Nevertheless, when used with the recommended setup, AInsectID Version 1.1 Color Merge excels in image segmentation and provides consistent performance for general applications, making it a valuable tool for various imaging tasks.

## Supporting information

read me

## Data Availability

All data set data is available as electronic supplementary material (ESM). The ESM include: (1) an .xlxs file including all data set data and (2) a .pdf READ ME file describing the contents of the .xlxs file.

## Author Contributions

Conceptualization (PA); Methodology (HS, PA); Software (HS, PA); Validation (HS, PA); Formal analysis (HS, SSM); Investigation (HS, SSM); Resources (PA); Data Curation (HS, SSM); Writing - Original Draft (HS); Writing - Review and Editing (PA); Visualization (HS, PA); Supervision (PA); Project administration (PA); Funding acquisition (HS)

## Acknowledgements

We wish to thank the Higher Education Commission (HEC) of Pakistan for funding the PhD scholarship of HS. We are also grateful to Dr. Marcelo Dias for the insightful discussions leading up to the submission of this article.

## A Standardized Setup

Standardized imaging setup for analyses conducted using the AInsectID Version 1.1 Color Merge algorithm. The setup includes four white LED lights, which should be positioned our corner arrangement, and at 45°angles relative to the specimen plane (cf. Figure 2). This ensures a balanced illumination is achieved, where the effects of individual light sources cancel each other out. A white background light (using a light board with controllable lux), is also applied, maintaining ambient lighting within a controlled lux range of 550–650. The camera should be positioned 10 cm above the specimen using a 1:1 magnification, with the camera oriented in the plan view relative to the specimen (the real image surface), using a 52 mm lens and an aperture of f/5.6 without digital zoom. In a plan (top-down) view, the camera is fixed at 90° to the center of the specimen plane, with no tilt. However, it may be rotated 360 ° around its vertical axis to adjust orientation while maintaining the consistent viewing angle. Original capture resolution should be at least 2048×2048 px (4 MP).

**Figure A1.**
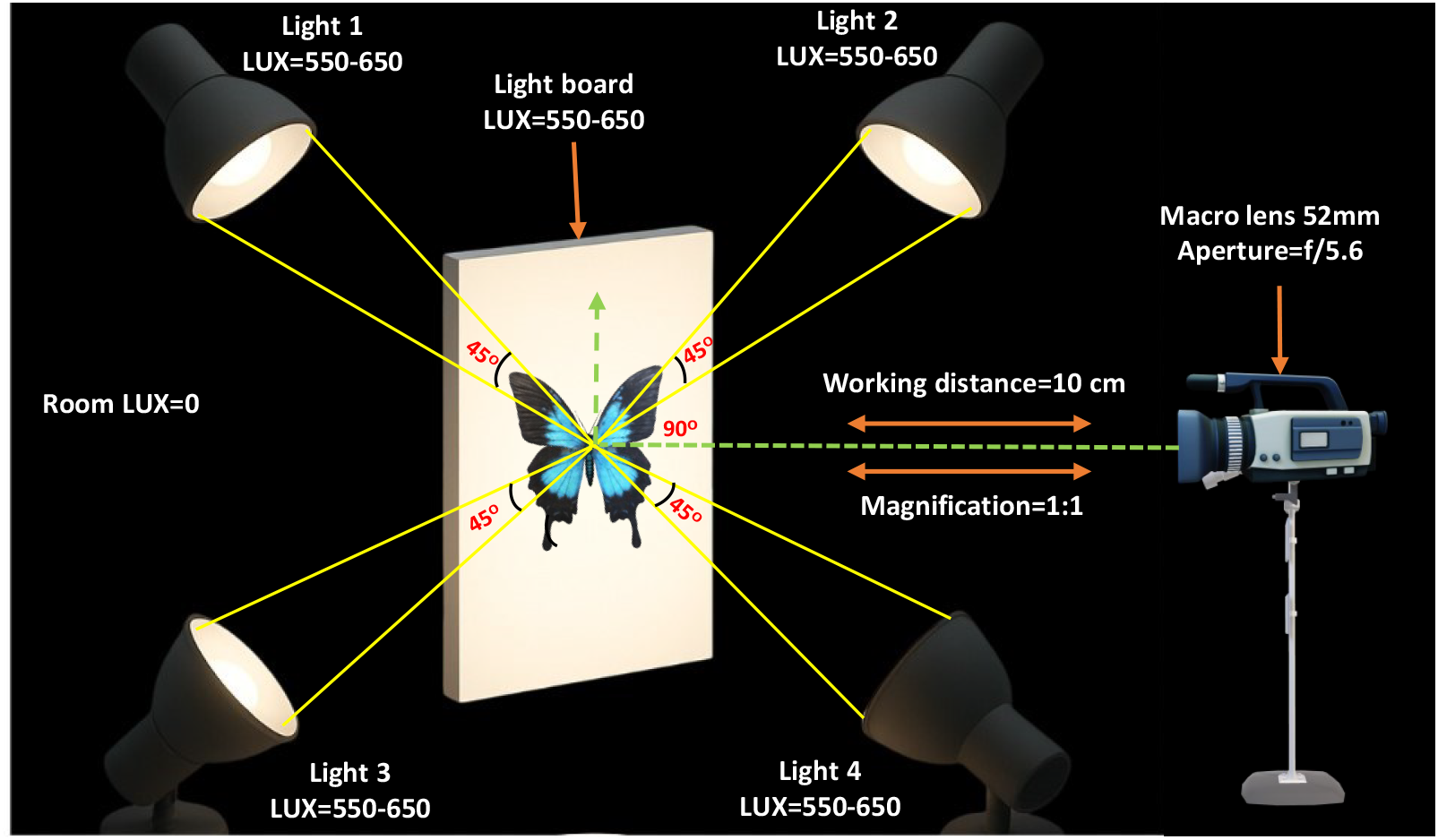
Schematic of the standardized imaging setup developed for use with the AInsectID Color Merge algorithm

