## Supplementary material for "A Standardized Method for Insect Color Analyses using Open Source Software: AInsectID Version 1.1 Color Merge": read me

### READ ME file on electronic supplementary data available in the .xlxs spreadsheets

Haleema Sadia[1],Sahal Sabilil Muttaqin[2], Parvez Alam[1*]
[1]School of Engineering, The University of Edinburgh, Edinburgh, UK
[2]School of Biological Sciences, The University of Edinburgh, Edinburgh, UK

May 28, 2025

#### 1 Introduction

The data set was created to evaluate and standardize the AInsectID Version 1.1 Color Merge algorithm, focused on a hybrid algorithm that integrates Simple Linear Iterative Clustering (SLIC), a computer vision algorithm used for the initial generation of superpixels, with the Delta E 2000 function to enable precise merging of superpixels based on perceptual color differences. The software outperforms previous segmentation approaches, achieving a Silhouette Score of 97%, which indicates high accuracy in distinguishing distinct color clusters [1–3]. Despite this improvement, the method remains sensitive to lighting conditions and the working environment, which can impact color consistency and segmentation reliability. To understand how different lighting conditions and working environments affect the performance of the AInsectID Version 1.1 Color Merge algorithm, we conducted a series of experiments and created this dataset. It was created by using butterfly wing color data of five different species under controlled imaging conditions:

- *Papilio glaucus*
- *Philolaus*
- *Battus philenor*
- *Diaethria*
- *Agrias amydon*

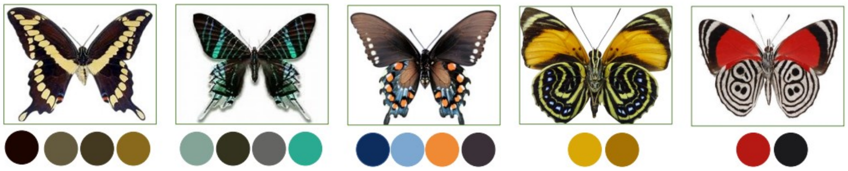

Figure 1: Color selection process: Five butterfly specimens were chosen, distinct colors are selected from each specimen for processing and analysis using the AInsectID Version 1.1 Color Merge algorithm

We analyze the color consistency of 16 distinct wing colors from five species of butterflies (see Figure 1). To understand how imaging parameters affect the performance of AInsectID Version 1.1 Color Merge, our experimental setup consists of:

- Illumination Intensity: ranging from 100 to 650 Lux with intervals of 50
- Number of Light Sources: 1 light, 2 lights, 3 lights, and 4 lights
- Working Distances: 10 cm, 20 cm, and 30 cm
- Camera Settings: SLR camera Monitech4K UltraHD 48MP, featuring a 16x digital optical zoom lens with 52mm focal length, f/5.0 aperture, and wide-angle lens (5.04mm), and 1:1 magnification.
- Color Channels Measured: Red (R), Green (G), and Blue (B)

We captured five separate images of each wing for each combination of settings (e.g., 100 lux, 10 cm distance, and one light source). These images were then processed using the AInsectID Version 1.1 Color Merge algorithm, and the RGB values for each color patch were extracted. We calculated the average RGB values across the five processed images for each color patch to ensure accuracy and reduce the effect of potential measurement fluctuations. This averaged RGB value was used as the final output for that specific setting DeltaE 2000 ($\Delta E_{00}$) is a color-difference metric that quantifies how perceptible the difference is between two colors, incorporating perceptual uniformnesses based on human vision. **Just Noticeable Difference (JND)** refers to the minimum $\Delta E_{00}$ value—typically around 1.0—at which a color difference becomes perceptible to the average human observer [3].

Our dataset consists of five spreadsheets in an .xlxs file (*Papilie Glaucuc*, *Philolaus*, *Battus philenor*, *Diaethris*, and *Agrias Amydon*), with each sheet representing the color data collected from an individual butterfly wing. This organization enables separate analysis and comparison across different wing samples.

### 2 Data Table Format

Each Excel sheet consists of four main data tables:

- **Color Block**
- **Delta E 2000 Color Difference**
- **p-value Analysis for RGB Channels**
- **Summary Table of p-values**

#### 2.1 Color Block

This table displays the RGB values of the colors in each lighting configuration, including variations in the lux level, number of light sources, and working distances. Each color block in the dataset corresponds to a specific working distance (10 cm, 20 cm, and 30 cm). Within each block:

- The table is divided into four groups, each representing the number of light sources used (from 1 light to 4 lights).
- For each light configuration, RGB values are recorded at increasing lux levels (100 to 650).
- **Example column headers:**
  - **Lux** – Illumination intensity
  - **R1, G1, B1** – RGB values under 1 light
  - **R2, G2, B2** – RGB values under 2 lights
  - **R3, G3, B3** – RGB values under 3 lights
  - **R4, G4, B4** – RGB values under 4 lights
- **Range** indicates how much the Red, Green, and Blue color values fluctuate as lux levels increase from 100 to 650. It reflects the sensitivity of each color channel to changes in lighting intensity under a fixed number of lights and working distance.

#### 2.2 DeltaE 2000 ($\Delta E_{00}$) color difference

This table presents the perceptual color differences $\Delta E_{00}$ between different lux levels under fixed lighting setups. A $\Delta E_{00}$ value greater than 1 indicates a perceivable difference to the human eye. The dataset is organized into four blocks, each corresponding to a fixed lighting setup with a working distance of 10 cm, 20 cm, and 30 cm and varying numbers of light sources (1 light, 2 lights, 3 lights, and 4 lights).

- Rows and columns represent the same set of lux levels (100, 150, . . . , 650).
- Each cell contains the $\Delta E_{00}$ color difference value between the color measured at the lux level of the row and the lux level of the column.
- The main diagonal (from top-left to bottom-right) is zero because it compares the same lux level with itself, so the color difference is zero.

**Sample Interpretation for Color 15 (Delta E 2000 color difference)**
**Condition:** Working Distance = 10 cm, 1 Light Source

The table entries show the $\Delta E_{00}$ color difference between image captures at different illumination levels (100 to 650 Lux). $\Delta E$ values greater than 1 indicate perceptible color differences to the human eye (above the Just Noticeable Difference threshold).

Several comparisons show $\Delta E_{00} > 2$, which suggests **noticeable or strong perceptual color shifts** due to varying illumination.

*Examples:*

- Between 100 Lux and 300 Lux: $\Delta E = 3.88$

- Between 150 Lux and 300 Lux: $\Delta E = 4.4138$ (very strong)
- Between 300 Lux and 400 Lux: $\Delta E = 4.6243$
- Between 100 Lux and 650 Lux: $\Delta E = 0.38$ (little or no visible change)

##### 2.3 P-value analysis for RGB channels

The table presents the P-values for the Red (R), Green (G), and Blue (B) channels under different lighting configurations—R1, R2, R3, and R4—measured at various lux levels ranging from 100 to 650 lux under fixed working distances (10 cm, 20 cm, and 30 cm). The goal is to evaluate whether the color channel intensities (R, G, B) show statistically significant differences when captured under different lighting configurations (e.g., R1 vs. R2, R1 vs. R3, etc.).

The P-value comparison is repeated for:

- **Red channel (R)**: 6 pairs — R1–R2, R1–R3, R1–R4, R2–R3, R2–R4, and R3–R4
- **Green channel (G)**: 6 pairs — G1–G2, G1–G3, G1–G4, G2–G3, G2–G4, and G3–G4
- **Blue channel (B)**: 6 pairs — B1–B2, B1–B3, B1–B4, B2–B3, B2–B4, and B3–B4

##### 2.4 Summary table of p-values for RGB channels

This table summarizes p-values for pairwise comparisons between different lighting setups (Light 1, Light 2, Light 3, Light 4) at three working distances (10 cm, 20 cm, and 30 cm), for two test samples, across all 16 colors.

The purpose of this study is to determine whether the lighting configuration significantly affects the color intensity in each RGB channel under different imaging distances and sample colors.

- **p-value $<$ 0.05** $\rightarrow$ Not statistically significant
- **p-value $\geq$ 0.05** $\rightarrow$ Statistically significant

**Sample Interpretation:** For *Color 15 at 10 cm working distance*:

- **Light 1–2:**
  - R: p = 0.388 $\rightarrow$ significant
  - G: p = 0.249 $\rightarrow$ significant
  - B: p = 0.824 $\rightarrow$ significant
- **Light 1–3:**
  - R: p = 0.0038 $\rightarrow$ No significant difference in red channel
  - G: p = 0.265 $\rightarrow$ significant
  - B: p = 0.857 $\rightarrow$ significant

This indicates that for Color 15 at 10 cm, the red channel is not sensitive to a change from Light 1 to Light 3, while the green and blue channels are significantly changed.

#### 3 Box-and-whisker Plot

The box-and-whisker plot is used to visually summarize the distribution of color difference $\Delta E_{00}$ data for each color sample. This plot helps to quickly identify central tendencies, data dispersion, and any anomalies within the dataset, making it an effective tool for comparing color stability across different lighting conditions (Lux: 100-650, number of lights) and working distances (10 cm, 20 cm and 30 cm). In the plot, the red boxes represent the color difference $\Delta E_{00}$ calculated at a working distance of 10 cm, the green boxes correspond to 20 cm, and the yellow boxes indicate 30 cm.

#### 4 Choropleth Map

The choropleth map provides a detailed analysis of the lighting setup by illustrating the distribution of color differences ($\Delta E_{00}$) across various illumination levels (lux) for each color. Each lux level is depicted as a distinct region, shaded according to its corresponding $\Delta E_{00}$ value, enabling a clear visual representation of how color variation changes with lighting intensity. This map helps identify the specific lux intervals within the optimal working distance and lighting conditions where $\Delta E_{00}$ values remain below the Just Noticeable Difference (JND) threshold $\Delta E_{00}$ less then or equal to 1), indicating color differences that are imperceptible to the human eye. Such visualization aids in pinpointing the lighting conditions that ensure stable and consistent color merging performance.

#### 5 Summary

This dataset has been carefully curated to support a thorough analysis of how different lighting conditions and working environments affect the performance of the AInsectID Version 1.1 Color Merge algorithm. By providing detailed measurements across multiple variables—including illumination intensity, number of light sources, working distances, and color channels—this dataset aims to facilitate reproducible and comprehensive evaluations.

Researchers and developers can use this dataset to better understand the algorithm's behavior under varying experimental setups, identify optimal configurations for accurate color merging, and further improve robustness against environmental factors.

[2] Haleema Sadia and Parvez Alam. AInsectID Version 1.1: an insect species identification software based on the transfer learning of deep convolutional neural networks. *Advanced Intelligent Discovery*, 2024. 202400032.

[3] Haleema Sadia and Parvez Alam. Enhanced K-means color clustering based on SLIC superpixels merging incorporated within the entomology software: AInsectID. In *2024 IEEE International Conference on Evolving and Adaptive Intelligent Systems (EAIS)*, pages 1–9. IEEE, 2024.
